# The evolution of family reputation extends indirect reciprocity

**DOI:** 10.64898/2026.08.27.747476

**Authors:** Miguel Dos Santos, Hisashi Ohtsuki, Charles Mullon

## Abstract

Reputation plays a major role in supporting cooperation among unrelated individuals through indirect reciprocity. By helping others, individuals build a good personal reputation and receive greater benefits from future partners. Most models of indirect reciprocity assume that a person’s reputation reflects only their own behaviour. Yet in many societies, people are also judged by their family’s reputation. How family reputation affects the evolution of cooperation, and whether reliance on it can itself evolve, remain unclear. Here we show that reputation inheritance expands the conditions under which indirect reciprocity favours cooperation, increasing helping and favouring greater reciprocity. Greater reciprocity in turn favours stronger reliance on inherited reputation, creating a positive feedback that stabilises cooperation, especially when interactions are infrequent or personal behaviour is difficult to observe. This feedback arises because cooperation generates future benefits both for the individual, through their personal reputation, and for their descendants, through inherited reputation. Reputation inheritance thereby provides a route via which kin selection and reciprocity, often treated as alternative explanations for cooperation, can reinforce one another. Our model helps explain why family-based reputation occurs across diverse human societies and provides an evolutionary framework for studying phenomena organised around family standing, including kin-based institutions, feuds between families and honour-based violence within them.

## Introduction

Reputation plays a key role in supporting human cooperation among unrelated individuals [1–9]. When helping is observed, it can build a personal reputation that brings subsequent benefits from third parties; a mechanism known as indirect reciprocity [1, 4]. Evolutionary models show that indirect reciprocity can maintain cooperation [4, 10–18], while experimental and field studies show that individuals with good reputations are more likely to receive rewards, trust and social support [2, 6, 9, 19–24].

Across diverse societies, however, individuals can also be judged by their family’s reputation, with consequences for trust, cooperation and access to social opportunities. We bring together published examples of this phenomenon in Table 1. More broadly, kin-based institutions have long organised economic and social life through systems of descent, marriage and residence [25]. Family reputation can operate in both directions: individuals may be evaluated partly through the reputation of their parents or wider family [26–31], while their behaviour may affect how other family members are perceived [32–34]. To the extent that behavioural tendencies and social values are transmitted within families [35, 36], family reputation may provide a cue to an individual’s likely behaviour before their own behaviour has been observed.

**Table 1.** Examples of the importance of family reputation across societies.

| **Mechanism** | **Examples across societies** |
| --- | --- |
| Individual actions affect family reputation | China: individual behaviour affects family “face” (*mianzi*) [26]; Rural Vietnam: individual behaviour influences household reputation across generations [59–61]; Spain: insults to individuals threaten household reputation [62, 63]; New Zealand: Māori individuals may be seen as representing their ancestors and wider kin group [64]. |
| Honour norms and family standing | Turkey: protection of family honour (*namus*), especially female purity [65]; North India, Bangladesh, Pakistan and the Indian diaspora: *izzat* emphasises maintaining family reputation [32–34]; Afghanistan, Pakistan, India, Albania, and the Middle East and North Africa: honour norms are socially enforced, with violations leading to ostracism and affecting work, education and politics [56–58, 66–69]; United States: honour culture in some southern states links personal behaviour to family reputation [70, 71]. |
| Economic trust and market interactions | India: merchant families rely on reputation for business trust; strategic marriages strengthen family standing [72]; Rural Vietnam: household reputation improves access to credit [59]; United States: business families maintain financial and social reputation [31, 73]. |
| Marriage and mate choice | United Arab Emirates: family reputation is important in spouse selection [74]; 67 pre-industrial societies: family background strongly influences parental spouse choice [28]; India: merchant families use marriage strategically to enhance reputation [72]. |
| Collective security and conflict | Pakistan (Marri Baluch): family reputation deters theft and kidnapping [25]; Venezuela and Brazil (Yanomamö): kin groups with reputations for retaliation are less likely to be attacked [55]. |
| Cooperation and social capital | Peru (Quechua): cooperative households develop stronger support networks and better health outcomes; household heads shape family reputation [29, 75]; Native North Americans: cooperative households gain political influence and trade advantages; potlatch ceremonies build family prestige [30, 76]; Japan and Korea: family reputation shapes community standing and gift exchange between family groups [77–79]. |
| Familism and collective identity | Latinos in the United States: familism emphasises protecting family reputation [27, 80]; Nigeria: familism stresses sustaining family reputation [81]. |

Despite its apparent importance in many societies, family reputation has received limited attention in models of social evolution. Most models track personal reputation [4, 10–18], whereas related work considers reputations assigned to social groups rather than transmitted among relatives [37–40]. One previous model showed that inheriting a parent’s binary reputation as an offspring’s initial reputation can stabilise conditional cooperation against opportunistic defection [41]. It therefore remains unclear under what conditions family reputation promotes indirect reciprocity, when reliance on family information itself evolves, and how cooperation is affected when reputational consequences are shared among relatives.

To address these questions, we develop an evolutionary model in which cooperative behaviour, reciprocity and reliance on family reputation can coevolve. We consider a population whose members each participate in *T* pairwise donation games with randomly selected partners (Supplementary Information A for details). In each interaction, a donation of *d* costs the donor *dc* and provides the recipient a benefit *db*, where *c* and *b* are the cost and benefit per unit donated. Payoffs are accumulated across interactions and determine reproductive success.

The amount that individuals donate depends on their baseline tendency to cooperate and on the reputation of their partner (*ρ*). Donation is determined by two traits: a baseline donation *α* ∈ [0, 1] and a level of reciprocity *β* ∈ [0, 1), which measures responsiveness to the partner’s reputation, such that the donation from individual *i* to partner *j* is *d*_*i*_ = *α*_*i*_ + *β*_*i*_ *ρ*_*j*_ . Larger values of *β* therefore correspond to stronger conditional cooperation. An individual’s personal reputation is publicly observable and reflects their past behaviour; specifically, personal reputation equals the donation made in the previous interaction as in public image scoring models [11, 12, 15]. At the beginning of each generation, however, individuals have not yet acted, and standard models assign them a default reputation [12, 14, 42]. Following [14], we set this default reputation as zero (sometimes referred to as a discriminator “prejudice” of 0 [42]) but later consider random default reputations [11, 42] and introduce errors in action execution and reputation assessment [13].

Our key extension is to allow individuals to use family reputation rather than relying only on this default evaluation. Before partner *j* ‘s first observed donation, individual *i* evaluates *j* using *ρ*_*j*_ = *h*_*i*_ *ρ*_F,*j*_, where *ρ*_F,*j*_ is the reputation attained by *j* ‘s parent at the end of the parent’s life (under image scoring, the parent’s final donation), and *h*_*i*_ is the weight that *i* places on this inherited information (Fig. 1). Thus, *h*_*i*_ = 0 recovers the conventional assumption that an unobserved individual has the default reputation of zero, whereas larger values of *h*_*i*_ imply greater reliance on family reputation. Once *j* ‘s own donation has been publicly observed, this inherited reputation is replaced by *j* ‘s personal reputation. We assume that parent–offspring relationships and reputations are public, and that all offspring of an individual inherit the same reputation. Reputation inheritance is therefore strictly from parent to offspring in the baseline model. We later consider a different system in which all family members share a common family reputation that remains available to others throughout life, as observed in some societies (Table 1). Finally, the three traits *α, β*, and *h* can all evolve and are transmitted vertically from parent to offspring, genetically or culturally. In our simulations, each trait can mutate with probability *μ* and mutational variance *σ*.

**Figure 1.**
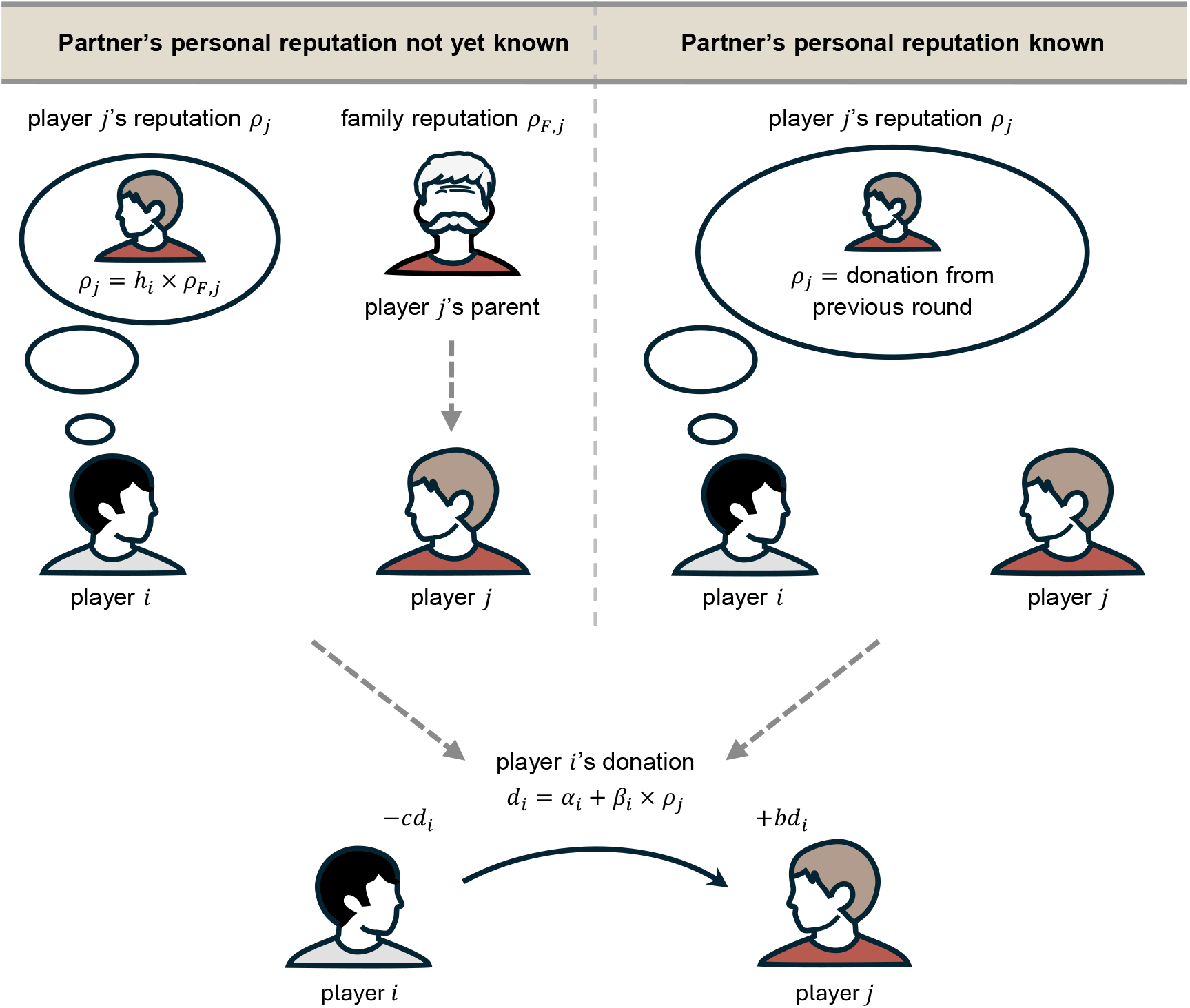
Family reputation in indirect reciprocity. An individual’s personal reputation *ρ* is their donation in the previous round (assuming public image scoring). If the partner has not yet been observed (e.g., in the first interaction at the beginning of life), a player may use the partner’s parental reputation (*ρ*_F_) to assess reputation, weighted by the extent to which the player relies on family reputation *h*. The donation *d* is determined by a baseline donation (*α*) and the responsiveness (*β*) to the partner’s reputation. The player then pays an amount *dc* for the partner to receive *db*, where *c* and *b* are the per-unit cost and benefit of cooperation. See Supplementary Information A.1 for more information on the model.

## Results and Discussion

### Family reputation favours cooperation

We first consider a baseline scenario in which reliance on family reputation (*h*) is fixed in the population (Supplementary Information B.1–B.2 for details). When *h* = 0, our model reduces to classical indirect reciprocity based exclusively on personal reputation [13]. In this case, selection favours greater baseline donation (*α*) and stronger reciprocity (*β*) when the resident level of reciprocity exceeds a critical threshold *β*^*^ (Fig. 2; Supplementary Fig. 1; Supplementary Information B.1, eqs. 27–28; [11, 14, 43]). Above this threshold (*β β*^*^), selection drives further increases in *α* and *β*, leading to a fully cooperative state in which both *α* and *β* evolve towards 1 (Fig. 2a; Supplementary Fig. 1a–c). The critical reciprocity threshold *β*^*^ therefore delimits the basin of attraction of the cooperative state: lowering *β*^*^ expands the range of initial conditions from which cooperation evolves and increases its resistance to evolutionary collapse. Consistent with classical results, the initial level of reciprocity required (*β*^*^) decreases as the benefit-to-cost ratio *b*/*c* and the number of interactions *T* increase (Fig. 2c; [14]). In the limit of infinitely many interactions (*T* → ∞), *β*^*^ converges to a minimum value of *c*/*b* in this model (Fig. 2c; Supplementary Information B.1, eq. 31).

**Figure 2.**
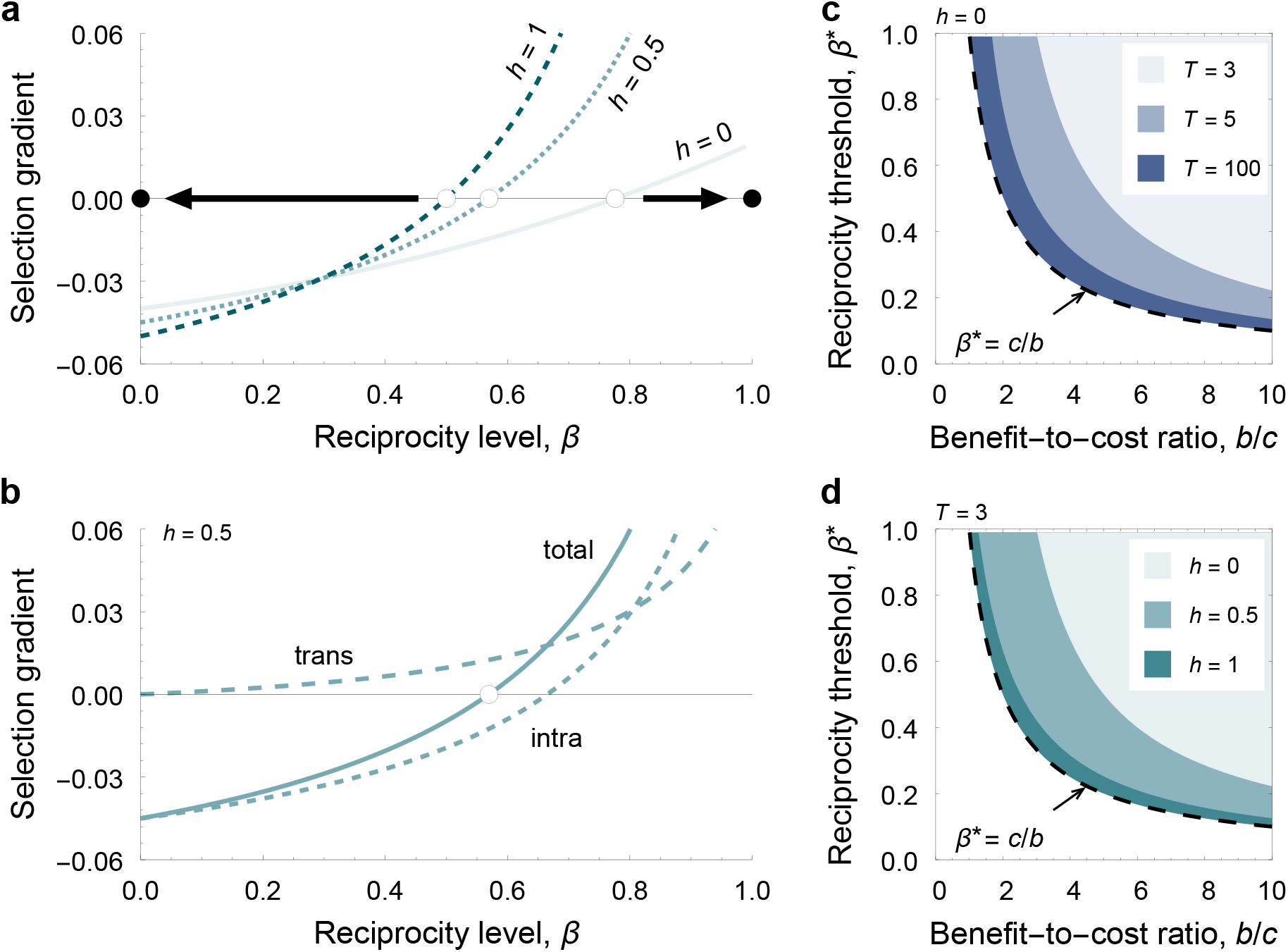
Reliance on family reputation favours cooperation. **a**, Selection gradient on reciprocity *β* indicating the direction of evolution favoured by selection: *S (β)* >0 and *S*(*β*) < 0 indicate selection for larger and smaller values of *β*, respectively, shown for different fixed levels of reliance on family reputation *h* in the population (Supplementary Information B, eq. 25b). Greater reliance on family reputation thus expands the basin of attraction for cooperation, by lowering the critical reciprocity threshold *β*^*^ required for further increases in *β* to be favoured. White and black dots denote unstable and stable equilibria, respectively. **b**, Decomposition of the selection gradient *S (β)* into intra- and transgenerational effects, which capture the effects of the reciprocity trait on the fitness of the individual and of all its descendants, respectively (Supplementary Information B, eq. 24). The cost of building a good personal reputation and the reciprocated benefits received within an individual’s lifetime thus often disfavours reciprocity (negative intragenerational effects). By contrast, inherited family reputation creates kin selection benefits that always favour reciprocity (positive transgenerational effects). **c, d**, Critical reciprocity threshold *β*^*^ above which higher reciprocity is favoured, shown as a function of the benefit-to-cost ratio (*b*/*c*). The threshold *β*^*^ is found by solving the zeroes of eq. 25b for *β*. **c**, Increasing the number of rounds *T* decreases *β*^*^, expanding the region where cooperation is stable. As *T* becomes large (e.g., *T* = 100), *β*^*^ approaches *c*/*b*. **d**, Greater reliance on family reputation similarly decreases *β*^*^, especially when *T* is small. As *h* approaches 1, *β*^*^ also approaches *c*/*b*, effectively extending the interaction horizon. Parameters: all panels, *c* = 1; **a–b**: *b* = 2, *T* = 5.

Reliance on family reputation expands the basin of attraction of cooperation (Fig. 2; Supplementary Fig. 1). Increasing *h* lowers the critical reciprocity threshold *β*^*^ required for selection to favour further increases in *β* (Fig. 2; Supplementary Fig. 1; Supplementary Information, eq. 41). Family reputation is used before a partner has established a personal reputation and therefore directly affects the initial interaction. Hence, the relative contribution of family reputation declines as personal observations accumulate, making its effect strongest when individuals interact only a few times (Supplementary Fig. 1d–f). When reliance on family reputation is complete (*h* = 1), the critical reciprocity threshold falls to *β*^*^= *c*/*b*, matching the limit obtained with infinitely many within-generation interactions (Fig. 2d and Supplementary Fig. 1c–f; Supplementary Information B.2, eq. 45). Thus, inherited reputation can sustain selection for reciprocal cooperation even when individuals interact only once in their lifetime (*T* = 1), provided that inherited information is used (*h* > 0).

Selection on reciprocity can be decomposed into intra- and transgenerational components (Fig.2b; Supplementary Information B, eq. 25b). The intragenerational component captures how responding to a partner’s reputation affects an individual’s payoffs and reputation during its own lifetime. The transgenerational component captures the resulting benefits to descendants that inherit the individual’s reputation. Because descendants of more reciprocal individuals inherit a better reputation and receive more help, this component always favours greater reciprocity and lowers the critical threshold (Fig.2b). Behaviour towards unrelated partners can therefore benefit descendants, extending indirect reciprocity beyond the individual’s lifetime and linking it to kin selection.

### Selection favours reliance on family reputation

Next, we allow reliance on family reputation *h* to evolve (Supplementary Information B.3). We find that selection favours higher *h* when baseline donation is positive (*α* > 0), and reciprocity in the population *β* is greater than *c*/*b* (Supplementary Information B.3, eq. 46) – a threshold lower than *β*^*^ (Supplementary Information B.3, eq. 48). Thus, reliance on inherited reputation can evolve once cooperation and sufficient reciprocity are present (Fig. 3a). Greater reliance increases an individual’s first donation, improving their personal reputation and, when reciprocated, their returns in later interactions. This intragenerational benefit allows *h* to evolve from zero. Once *h* > 0, parental reputation also benefits descendants before their own behaviour is observed, strengthening selection for reciprocity. Greater reciprocity, in turn, favours higher *h*, creating a positive evolutionary feedback (Fig. 3a).

**Figure 3.**
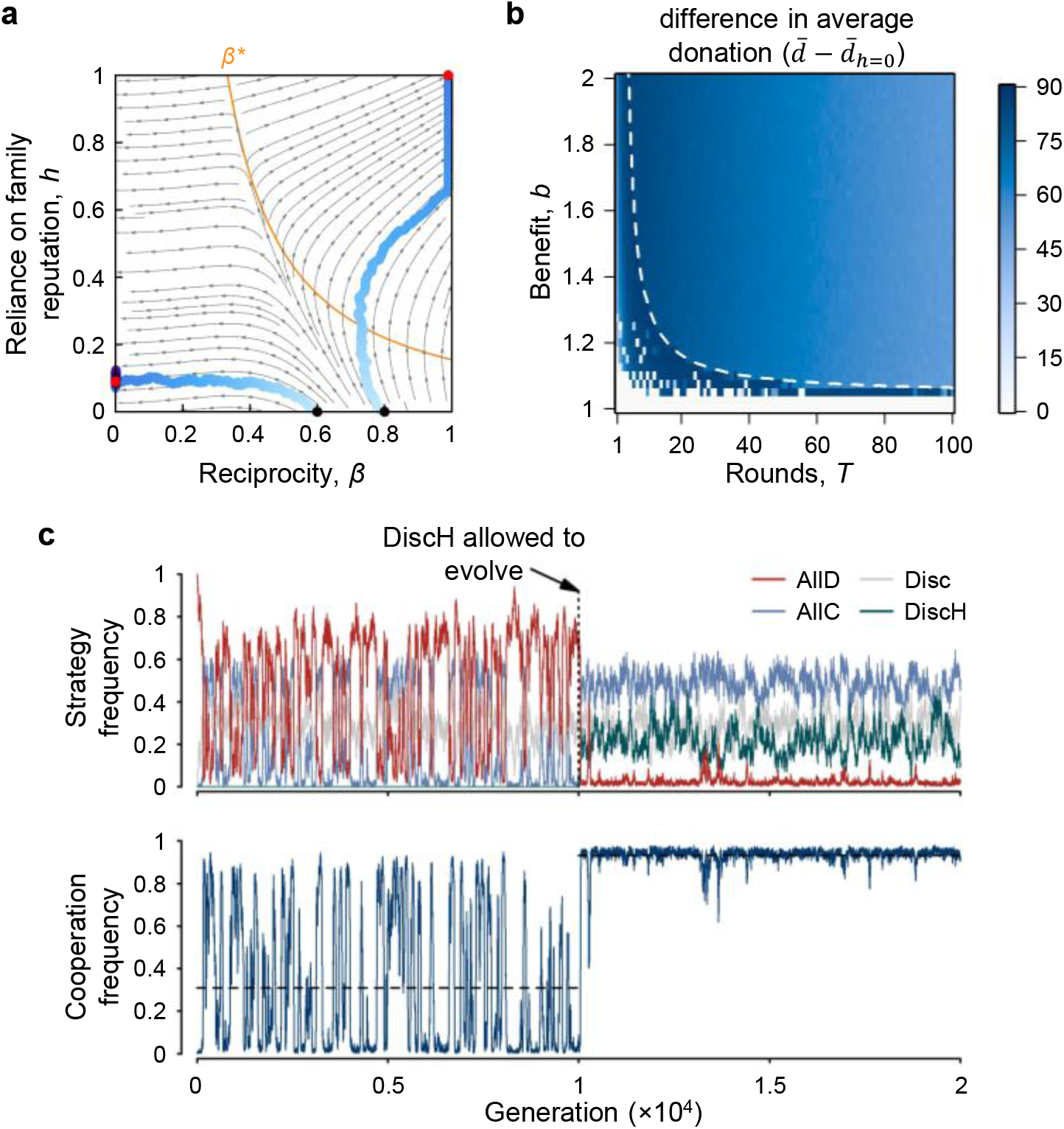
Joint evolution of cooperation and reliance on family reputation. **a**, Joint evolutionary dynamics of reciprocity *β* and reliance on family reputation *h* (Supplementary Information, eq. 61), when baseline donations are fixed in the population (*α* 0.1). Blue dots show mean trait values from stochastic simulations, with darker shades indicating later generations, and with black and red dots indicating initial and final trait values, respectively (Supplementary Information C for details on simulations). The orange line shows the critical reciprocity threshold *β*^*^, above which greater reciprocity is favoured. Reciprocity initially declines below this threshold but selection simultaneously increases *h* which lowers *β*^*^. The population can thus cross this boundary, after which *β* and *h* increase together through positive feedback. **b**, Difference in average donations between simulations in which reliance on family reputation *h* was and was not allowed to evolve (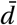 and 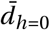, respectively, measured over the final third of generations), as a function of the number of rounds *T* and benefit *b*. Allowing *h* to evolve both expands the parameter region in which cooperation is stable, and substantially increases donations. The dashed line shows the region above which cooperation is stable without family reputation. **c**, Strategy and cooperation frequency in a model with pure strategies (Supplementary Information C.4 for details on simulations). Without family reputation, cooperation is unstable: discriminators (Disc) suppress unconditional defectors (ALLD), but unconditional cooperators (ALLC) can then replace Disc through drift and are subsequently invaded by ALLD. Allowing discriminators who rely on family reputation (DiscH) to evolve substantially stabilises cooperation. Parameters: all panels, *c* = 1, *q* = *s* = 1, ϵ_e_ = ϵ_a_ = 0; panel **a**: *N* = 10^4^, *T* = 2, *b* = 3, *ρ*_ini_ = 0, *µ* = *σ* = 10^−3^; panel **b**: *N* = 10^3^, *ρ*_ini_ = 0, *µ* = 10^−3^, *σ* = 0.1; panel **c**: *N* = 10^3^, *T* = 20, *b* = 3, *ρ*_ini_ = BAD, *µ* = 0.01.

Individual-based simulations confirm that reliance on family reputation evolves across a broad range of conditions and stabilises cooperation where it would otherwise collapse (Fig. 3b; Supplementary Fig. 2; see Supplementary Information C for details on simulations), especially when interactions are few and benefits are modest (low *T* and *b*/*c*). Where cooperation is already stable without family reputation, the evolution of *h* increases donation levels (Fig. 3b; Supplementary Fig. 2). These findings are robust to errors in action execution and reputation assessment (Supplementary Figs. 3–4), and when unknown partners are assigned random reputations (Supplementary Fig. 5).

Our analyses so far assume continuous traits and mutations of small phenotypic effect, whereas many models of indirect reciprocity consider a finite set of discrete strategies [11, 13, 15, 17, 40]. To test whether our conclusions depend on this assumption, we constructed a discrete-strategy model containing the three classical strategies: unconditional defectors, unconditional cooperators, and discriminators [14]; and a fourth strategy that behaves like a discriminator but also uses family reputation when evaluating an unobserved partner (Supplementary Information C.4). Previous work has shown that the image scoring norm generally cannot maintain full cooperation when family reputation is absent [11– 13, 15, 44]. Our results show that inheritance of family reputation not only evolves, but also increases and stabilises cooperation in this alternative framework (Fig. 3c). These results hold when actions and reputation assessments are subject to error (Supplementary Fig. 6) and individuals assume that unknown partners cooperate (Supplementary Fig. 7).

### Reliance on family reputation compensates for low observability

We now relax perfect observability of personal behaviour and perfect knowledge of family reputation (Supplementary Information C.2). Each interaction is now publicly observed with probability *q*, in which case personal reputations are updated; otherwise, they remain unchanged. Personal reputation may therefore remain unavailable for several interactions early in life (Supplementary Information C.2, eq. 64). If personal reputation is unavailable, family reputation is known with probability *s* and may be used; otherwise, partners assign the default reputation (zero).

Lower observability (low *q*) generally favours reliance on family reputation by increasing the opportunities to use it (Fig. 4a; Supplementary Fig. 8). Across most of the parameter range explored, reliance on family reputation again evolves and sustains cooperation where personal reputation alone cannot; the exceptions occur when reputations are rarely updated (*q* < 0.1) or family reputation is seldom known (*s* < 0.1; Fig. 4a; Supplementary Fig. 8).

**Figure 4.**
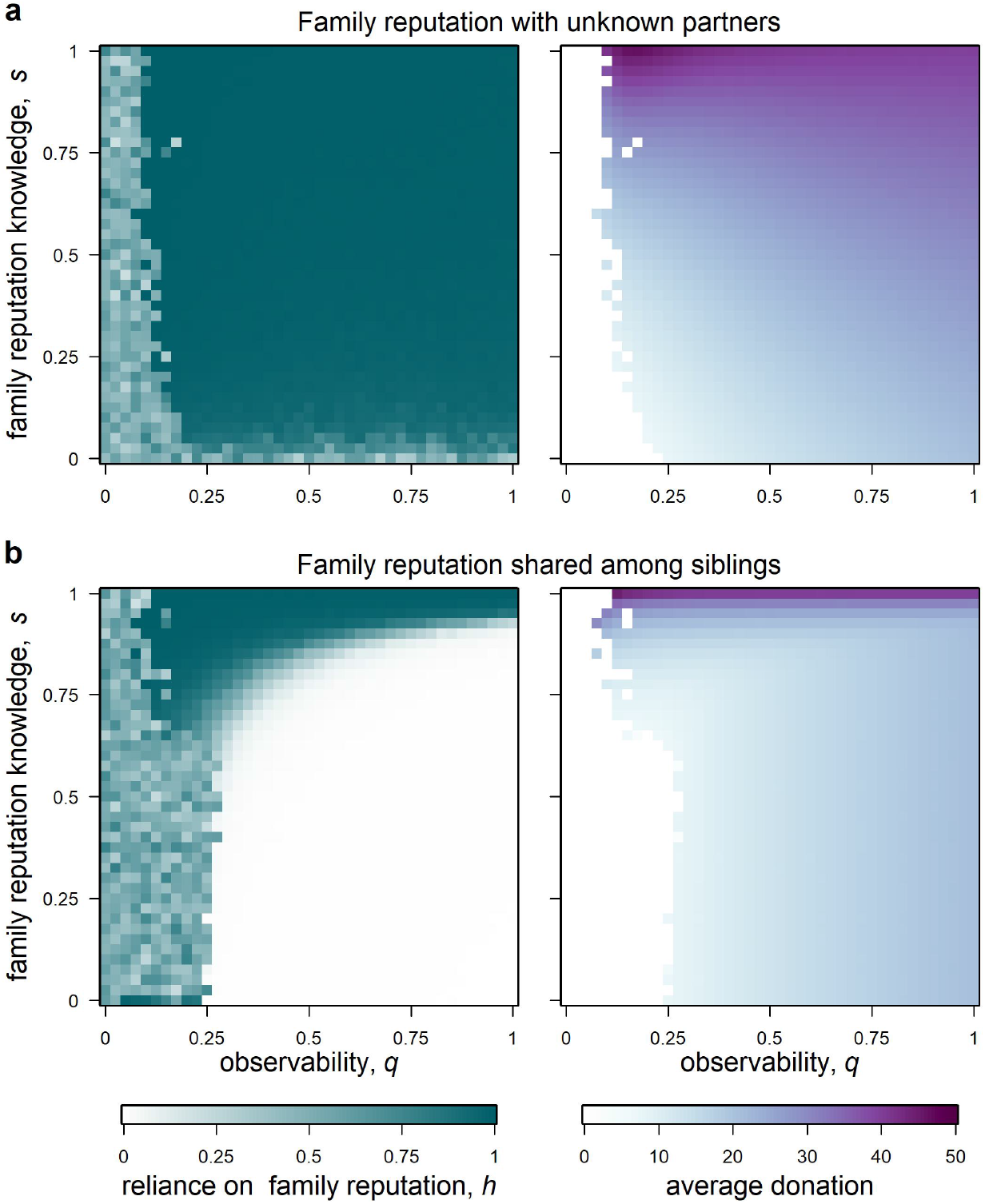
Family reputation compensates for low observability. Long-term average reliance on family reputation (*h*) and donation in stochastic simulations for different levels of interaction observability (*q*) and probabilities of knowing family reputation (*s*). **a**, Baseline scenario in which family reputation can be used only when the partner’s behaviour has never been observed (typically early in life, Supplementary Information C.2 for details on simulations). Reliance on family reputation evolves unless observability is very low (*q* < 0.15) and family reputation is rarely known (*s* < 0.1). When family reputation is well-known (*s* ≈ 1), lower observability increases donations, because family reputations accumulate over generations whereas personal reputations are updated less frequently. **b**, Siblings all share a single family reputation defined by the most recently publicly assessed donation made by any sibling, which can be used in every interaction (Supplementary Information C.3 for details). In this case, reliance on family reputation evolves when family reputations are sufficiently well-known (*s* > 0.6, and *q* > 0.25), because missing family information can now occur throughout life, rather than only before a partner is first observed. When family reputation is well-known (*s* > 0.6), reducing observability again favours reliance on family reputation. Unlike the baseline scenario, however, reliance on family reputation can be selected against, despite reciprocity being present (white area in the left panel). Parameters: *N* = 10^3^, *T* = 20, *c* = 1, *b* = 2, *ρ*_ini_ = 0, *µ* = 10^−3^, *σ* = 0.1, ϵ_e_ = ϵ_a_ = 0.05.

Where it evolves, reliance on family reputation also increases donations under imperfect observability (Fig. 4a; Supplementary Fig. 8). When observability is low and interactions are limited, personal reputations are updated too infrequently to reflect individual behaviour reliably within a lifetime (Supplementary Fig. 9). Inherited reputation prevents this information from being lost between generations. Because offspring also inherit behavioural tendencies from their parents, reputation can continue to track behaviour across generations and converge over time, even when it cannot do so within a single lifetime. This allows cooperative individuals to benefit earlier from indirect reciprocity (Supplementary Fig. 9). These effects persist in the presence of errors (Fig. 4a) and with discrete strategies (Supplementary Fig. 10).

### Reciprocity with shared family reputations

We have assumed that reputation is transmitted vertically, from an individual to their offspring. In many societies, however, one family member’s actions can affect how other relatives are judged by the community (Table 1), which amplifies the consequences of personal actions. To capture this, we now consider that siblings share a common family reputation, updated to the most recent publicly assessed donation made by any sibling (Supplementary Information C.3). An individual can therefore improve or harm the reputation of all siblings within a generation. For simplicity, individuals rely exclusively on either personal or family reputation, but not both; with *h* being the probability of using family reputation throughout life.

Reliance on family reputation can also evolve under this strong form of reputation sharing (Fig. 4b). As before, few interactions and low observability promote its evolution (low *T* and *q*, respectively), but family information must now be widely available (high *s*; Fig. 4b). If *s* is too low, reliance on family reputation is selected against even in fully cooperative populations (Fig.4b). This is because family reputation now replaces personal reputation throughout life: individuals who lack family information cannot evaluate their partners, and their resulting donations can reduce the shared reputation of their siblings. When family information is sufficiently available, however, *h* evolves, shared reputations increase and donations rise (Fig. 4b). Thus, the evolution of reliance on shared family reputation depends both on its availability and on whether it supplements or replaces personal information, potentially helping to explain variation in its use among societies.

### Conclusion

Overall, our results show that reliance on family reputation – a documented form of social evaluation across diverse human societies (Table 1) – can stabilise cooperation by expanding the conditions under which indirect reciprocity operates and can increase donations in cooperative populations. This occurs even when interactions are rare, behavioural information is scarce or errors in action and assessment are frequent: conditions under which indirect reciprocity based solely on personal reputation often breaks down [11, 13–15, 44]. Inherited reputation allows descendants whose behavioural tendencies resemble those of their parents to benefit from a good reputation even before their own behaviour is observed. Reputational consequences therefore cross generations, extending the effective horizon of reciprocity and coupling indirect reciprocity with kin selection. Reputation inheritance parallels other forms of non-genetic inheritance, such as ecological [45, 46], status [47] and wealth inheritance [48, 49].

Our results also show that reliance on family reputation is itself favoured by selection under a broad range of conditions, especially when personal behaviour is difficult to observe but family information remains available. By using inherited reputations, individuals can make better-informed decisions when interacting with previously unobserved partners, improving the effectiveness of indirect reciprocity. This increases the benefits of maintaining a good reputation for both the individual and their descendants. Our model does not consider the cognitive costs of remembering and updating reputations [39]. Replacing several personal reputations with a single family-level evaluation conceivably reduces these costs, at the expense of overlooking potential differences among family members. This shortcut should be most useful when family membership predicts behaviour well enough to compensate for this loss of individual detail. It may therefore make family reputation a practical basis for social evaluation in societies organised around kin-based institutions [25]. More broadly, assigning a common reputation to all family members may provide a cognitive basis for evaluating people according to group membership, contributing to stereotypes and collective reputations beyond the family [37, 39, 40, 50].

Our model isolates how reliance on family reputation evolves and affects cooperation in a simplified setting with public assessment and first-order social norms. The question now is how these dynamics unfold under private assessment [51], higher-order norms [52], and the complex group structures and sex-dependent inheritance patterns found in human societies [25, 53], and how these richer environments shape social behaviour within and between families. Beyond cooperation, family reputation may also govern costly retaliation and punishment where family honour is collectively defended. In feuds and vendettas, individuals may retaliate at personal cost to preserve their family’s honour or standing [54]. By building a collective reputation for vengefulness, such retaliation may deter future aggression, but it may also invite counter-retaliation and protract conflict [55, 56]. Perceived threats to family honour can likewise be invoked to justify honour-based abuse within families, from coercive control to lethal violence [57, 58]. Family reputation may therefore have opposing social consequences: supporting cooperation under some conditions while sustaining harmful norms and cycles of violence in others.

## Supplementary Information

## A. Baseline model

In this appendix, we present our baseline model in detail.

### A.1 Life-cycle, social interactions, and fitness

We consider a well-mixed population of large size *N* with the following life cycle: (i) first, adults engage in social interactions and obtain payoffs; (ii) they reproduce with fecundity determined by these payoffs; and (iii) they die, after which their offspring compete uniformly for recruitment into the next generation of adults. We detail these steps below.

#### A.1.1 A donation game

Individuals play a continuous donation game [1–4] over a finite number of rounds, *T*, interacting with a randomly selected partner in each round. In each round *t* ∈ {1, 2, …, *T*}, each individual independently chooses how much to donate to their partner. An individual’s donation depends on their traits and their partner’s reputation.

Three individual traits govern this donation rule: (i) the baseline donation, *α* ∈ [0, 1], which is independent of the partner’s reputation; (ii) reputation-based reciprocity *β* ∈ [0, 1), which determines the strength of the response to the partner’s reputation; and (iii) the tendency to use family reputation, *h* ∈ [0, 1], which determines the weight placed on the partner’s inherited reputation when that partner has not yet been observed and hence has the default personal reputation *ρ*_ini_ = 0.

Suppose that individuals *i* and *j* are paired at round *t* . The donation made by individual *i* is

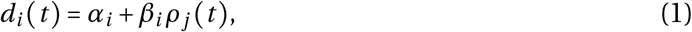

where *α*_*i*_ is individual *i* ‘s baseline donation, *β*_*i*_ is individual *i* ‘s reputation-based reciprocity, and *ρ*_*j*_ *t* is the reputation assigned to *j* by *i* at the beginning of round *t* . Similarly, the donation received by individual *i* in round *t* is

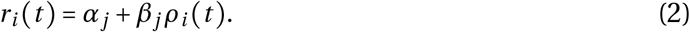

#### A.1.2 Reputation dynamics

We consider a simple first-order assessment norm analogous to image scoring [5], extended to allow reputations to be inherited. Specifically, the reputation that individual *i* assigns to its partner *j* in round *t* is

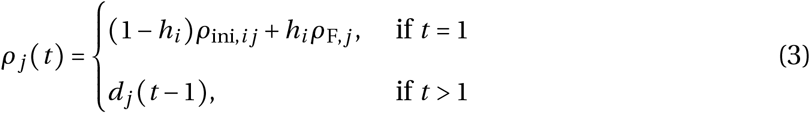

where *h*_*i*_ ∈ [0, 1] is individual *i* ‘s tendency to use inherited reputation, *ρ*_ini,*ij*_ is the default initial reputation that individual *i* assigns partner *j*, and *ρ*_F,*j*_ is the reputation inherited by individual *j*, equal to their parent’s final donation before reproduction. More precisely, if *p*(*j*) denotes the parent of *j*, then

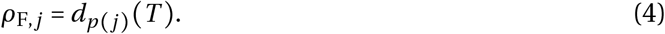

Thus, in the first round, *i* evaluates *j* using *j* ‘s inherited reputation, weighted by *h*_*i*_ . In all subsequent rounds, *i* evaluates *j* using *j* ‘s donation in the preceding round. In the baseline model, we assume that *ρ*_ini,*ij*_ = *ρ*_ini_ = 0, but we later allow individuals to be assigned a random initial reputation rather than zero. Because the first-round assessment depends on the observer through *h*_*i*_ and *ρ*_ini,*ij*_, it may differ among individuals. We suppress the observer index and write this assessment as *ρ*_*j*_ (*t*) to simplify notation.

When *h*_*i*_ = 0, individuals begin life with a reputation *ρ*_ini_ = 0 (as in e.g. [6]). When *h*_*i*_ > 0, by contrast, the reputation inherited from *j* ‘s parent affects the donation that *j* receives from individual *i* in the first round. Hence, even though we assume a public assessment system [7], individuals may still differ in how they assess a particular partner in the first interaction (reflecting a private reputation assessment; [8]). In later extensions, we allow inherited reputation to be used in any round in which personal information is not yet available.

#### A.1.3 Fitness

Donations incur a cost to the donor and provide a benefit to the recipient. We assume that these effects determine fecundity linearly such that the fecundity of individual *i* is

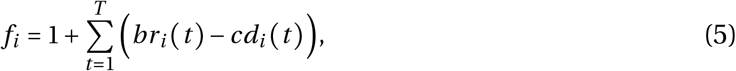

where *r*_*i*_ (*t*) is the donation received by *i* at round *t, d*_*i*_ (*t*) is the donation made by *i* at round *t*, and *c* > 0 and *b* > *c* denote the cost and benefit per unit donated, respectively. The constant 1 is baseline fecundity, and we assume that the parameters ensure *f*_*i*_ > 0 for every possible interaction history.

Because offspring compete uniformly for *N* positions in the next generation, the expected number of adult offspring produced by individual *i*, i.e. its fitness, is its fecundity relative to mean fecundity:

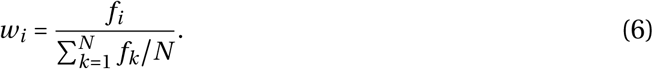

### A.2 Evolutionary dynamics

We are interested in the joint evolution of the three individual traits: *α, β*, and *h*. We assume that these traits are transmitted from parents to offspring genetically or culturally and that mutations or cultural innovations are rare and have small phenotypic effects. Simulations presented below show that our predictions are robust to changes of larger effect.

Under these assumptions, evolution can be studied by considering the invasion fitness of a rare mutant expressing trait values ***z***_m_ = (*α*_m_, *β*_m_, *h*_m_) in a resident population that is effectively monomorphic for trait values ***z*** = (*α, β, h*) [9–11]. Using eq. (6), invasion fitness in this model is given by

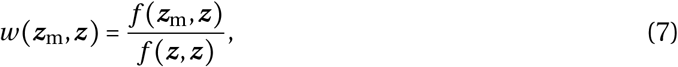

where *f* (***z***_m_, ***z***) is the fecundity of the mutant in the resident population and *f* (***z***, ***z***) is the fecundity of a resident individual in that population. Equation (7) assumes that *N* is sufficiently large for interactions between mutants to be negligible. We next derive the fecundities of the resident and of the mutant.

#### A.2.1 Resident fecundity

Consider a monomorphic resident population in which all individuals express phenotype ***z*** = (*α, β, h*). Because all individuals have the same phenotype, they make and receive the same donation in each round. Let *d*(*t*) denote this donation (with *r*(*t*) = *d*(*t*) and let *ρ*_F_ denote the reputation inherited from the previous generation.

In the first round, individuals evaluate their partners using the effective reputation *hρ*_F_. The initial donation is therefore

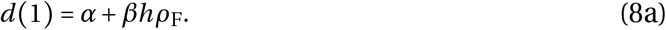

From the second round onward, reputation equals the donation made in the preceding round. Donations therefore follow the recursion

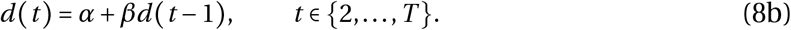

Solving this recursion (eq. 8b) with the initial condition eq. (8a) gives

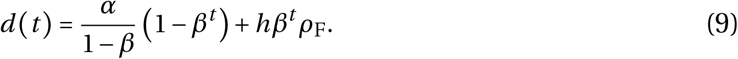

This allows us to characterise the reputation transmitted to the next generation, which recall is the donation made in the final round:

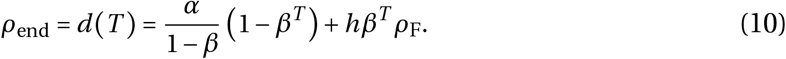

At equilibrium across generations, the inherited reputation equals the terminal reputation, so that 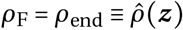. Solving this equilibrium condition using eq. (10) gives

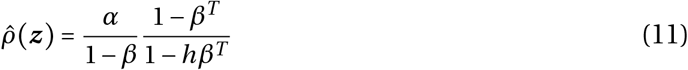

for the equilibrium family reputation in a resident population expressing ***z*** . Because *β* < 1, 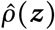 converges to *α*/(1 − *β*) as *T* → ∞.

The donation of a resident to a resident (eq. 9) can therefore be written as

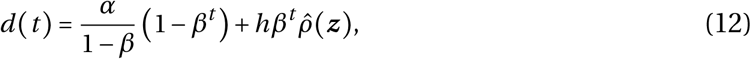

where 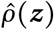 is given by eq. (11). Summing eq. (12) over the *T* rounds appropriately (as in eq. 5) then obtains

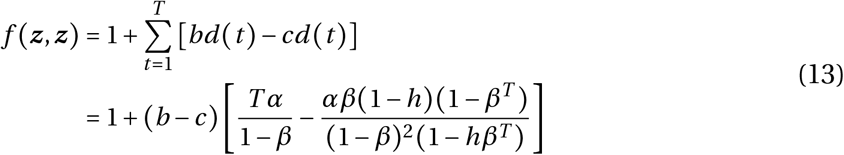

for resident fecundity.

#### A.2.2 Mutant fecundity

We now consider a lineage of rare mutants with phenotype ***z***_m_ = (*α*_m_, *β*_m_, *h*_m_) in a population that is monomorphic for the resident phenotype ***z*** = (*α, β, h*). To characterise the fecundity of a mutant (according to eq. 5), we need to track two donation trajectories: the donation *d*_m_ (*t*) made by a mutant to a resident, and the donation *d*_rm_ (*t*) made by a resident to a mutant (mutants are sufficiently rare that mutant–mutant interactions can be neglected).

In the first round, a mutant evaluates the resident using the resident’s equilibrium inherited reputation 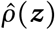, whereas the resident evaluates a mutant using the mutant’s inherited reputation, which we denote as *ρ*_F,m_. From the second round onward, each individual evaluates its partner using that partner’s donation in the preceding round. The donation trajectories therefore satisfy

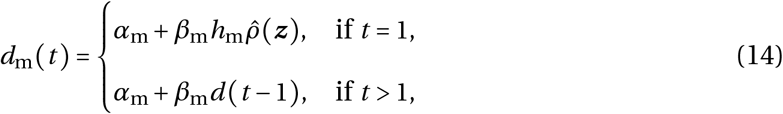

And

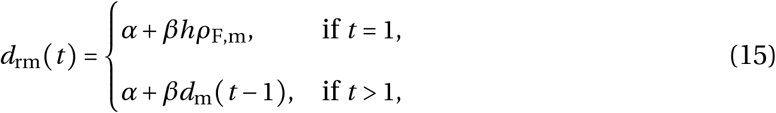

where 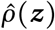 is given by eq. (11) and *d* (*t* − 1) is given by eq. (12).

Substituting eq. (12) into eq. (14) gives

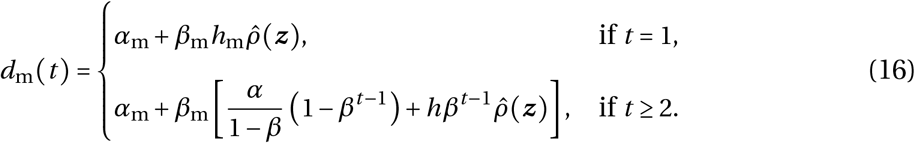

In turn, plugging eq. (16) into eq. (15) gives us

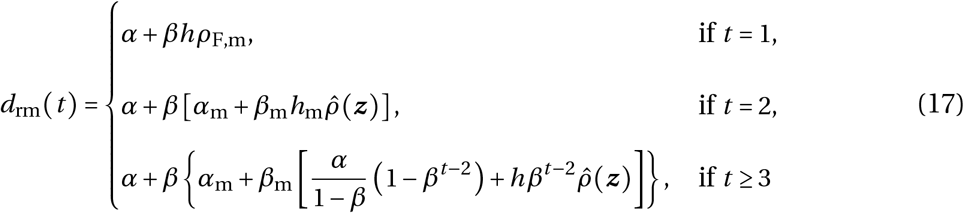

for the donation made by a resident to a mutant.

Let 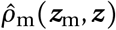 denote the equilibrium reputation inherited within a rare mutant lineage. This reputation equals the mutant’s donation in its final round and is given by

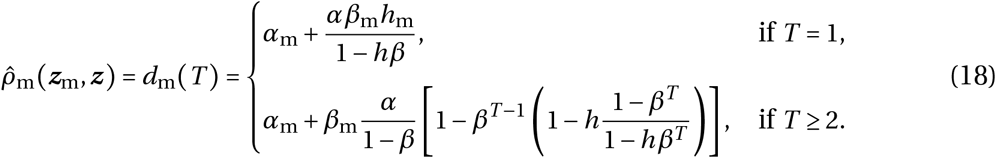

A newly arisen mutant initially inherits the reputation of its resident parent. Although this initial reputation affects the donation that the mutant receives in its first round, it does not affect the mutant’s own final donation, *d*_m_ (*T*). The reputation transmitted by the mutant is therefore determined entirely by its phenotype ***z***_m_ and the resident phenotype ***z*** . Because every mutant descendant makes the same final donation while the lineage remains rare, the inherited reputation within the mutant lineage reaches 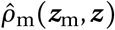 after one generation. As a consistency check, we have 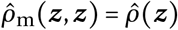, where 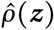is given by eq. (11). Because invasion fitness measures the asymptotic growth of the mutant lineage, we evaluate mutant fecundity at 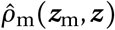 rather than at the reputation initially inherited from the mutant’s resident parent.

It will be useful later to distinguish the direct effect of a mutant trait from its effect through inherited reputation. To do this, we let Φ (***z***_m_, *ρ*_F,m_, ***z***) denote the fecundity of a mutant with phenotype ***z***_m_ and inherited reputation *ρ*_F,m_ in a resident population with phenotype ***z***, with *ρ*_F,m_ is as an independent argument. Following eq. (5),

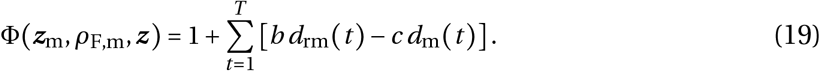

Substituting the donation trajectories from eqs. (16) and (17) and summing over rounds gives

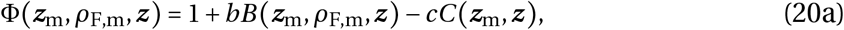

where *B(* ***z***_m_, *ρ*_F,m_, ***z)*** is the total donation received by the mutant and *C* ***z***_m_, ***z*** is the total donation made by the mutant. These are given by

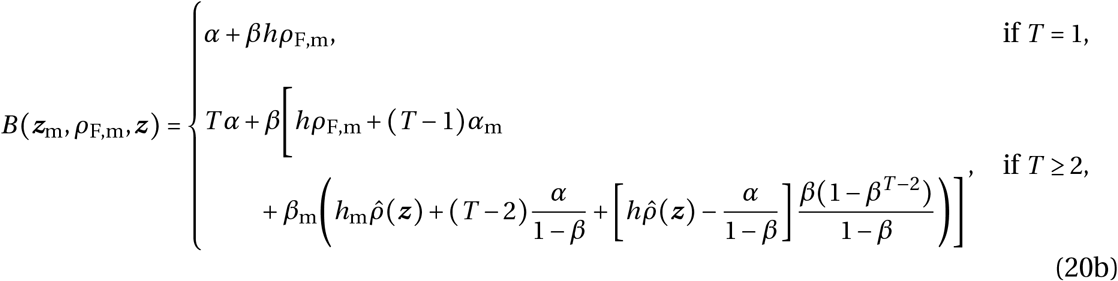

And

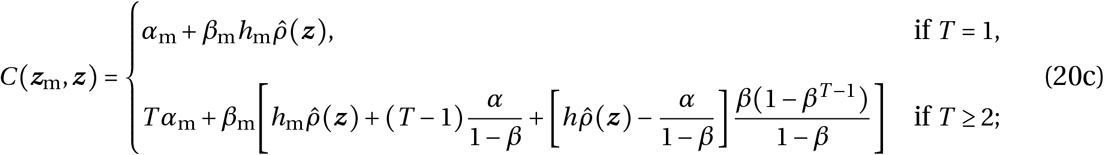

with 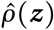 given by eq. (11).

The function Φ treats inherited reputation as independent of the mutant phenotype. Evaluating this reputation at its equilibrium value gives the fecundity of a mutant:

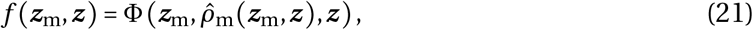

where the function Φ is given by eq. (20) and 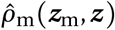 by eq. (18).

#### A.2.3 Selection gradient

Substituting eq. (21) into eq. (7), the fitness of a mutant lineage is

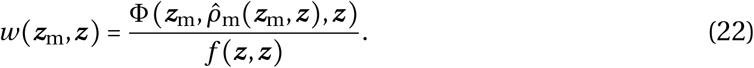

Assuming *N* is large enough such that mutant dynamics can be modelled as a branching process [12], a mutant lineage has a positive probability of invasion only if *w* (***z***_m_, ***z***) > 1. If *w* (***z***_m_, ***z***) ≤ 1, the mutant lineage eventually becomes extinct with probability one.

The selection gradient on each trait *a* ∈ {*α, β, h*} is given by

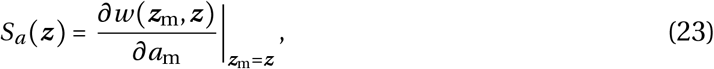

where *a*_m_ denote the value of trait *a* in the mutant. The selection gradient indicates the direction of evolution favoured by selection: *S*_*a*_(***z***) > 0 indicates selection for larger values of *a*, whereas *S*_*a*_ (***z***) < 0 indicates selection for smaller values in resident population expressing ***z*** . The three selection gradients form the vector ***S*** (***z***) = *S*_*α*_(***z***), *S*_*β*_ (***z***), *S*_*h*_ (***z***), which points in the direction favoured by selection in trait space. Evolution proceeds until it reaches a boundary of the trait space [0, 1] × [0, 1] × [0, 1] or an evolutionary singular point where all selection gradients vanish [9, 13].

The selection gradient in our model has two components because a mutant trait can affect fecundity through two routes. First, it can alter the mutant’s behaviour during its lifetime. Second, it can alter the reputation transmitted within the mutant lineage and thereby affect fecundity in subsequent generations. In this sense, inherited reputation acts as an extended phenotype [14], allowing selection to be decomposed into intragenerational and transgenerational effects [15, 16]. We obtain this decomposition by applying the chain rule to eq. (22):

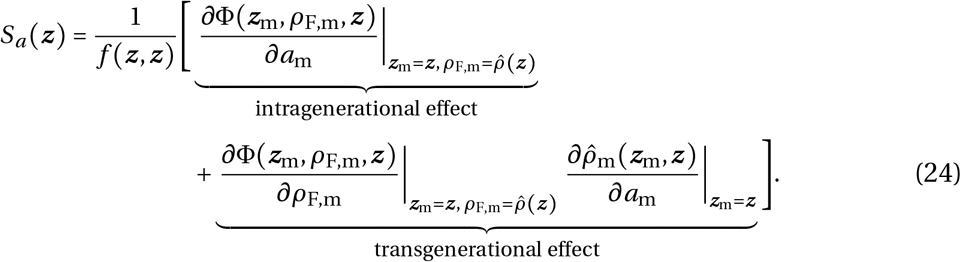

The intragenerational effect measures how a change in trait *a* affects mutant fecundity while inherited reputation is held constant. The transgenerational effect measures how the same change alters the equilibrium reputation inherited within the mutant lineage and how this altered reputation, in turn, affects mutant fecundity. This decomposition will be useful to understand the nature of selection later.

## B Analysis of selection and evolutionary dynamics

We now analyse selection on each of the three traits and derive the mathematical results underlying the main text.

Under weak mutation, the evolutionary change in each trait is proportional to its selection gradient [9]. Thus, for each *a* ∈ {*α, β, h*}, Δ*a* ∝ *S*_*a*_ (***z***), where the positive proportionality factor depends on the mutation rate and mutational variance. Substituting eqs. (18) and (20) into eq. (24) gives the three selection gradients. Each gradient contains the common positive factor 1 / *f* (***z***, ***z***). Because this factor affects only the rate of evolution, we absorb it, together with the mutation-related factors, into the proportionality constant. Up to this positive rescaling, the joint evolutionary dynamics of all three traits are therefore

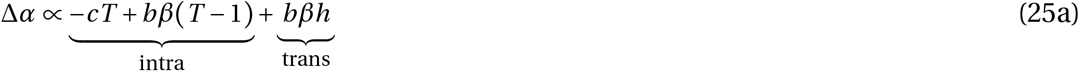

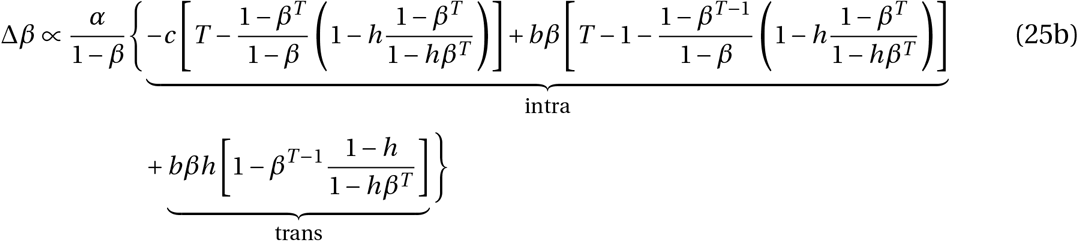

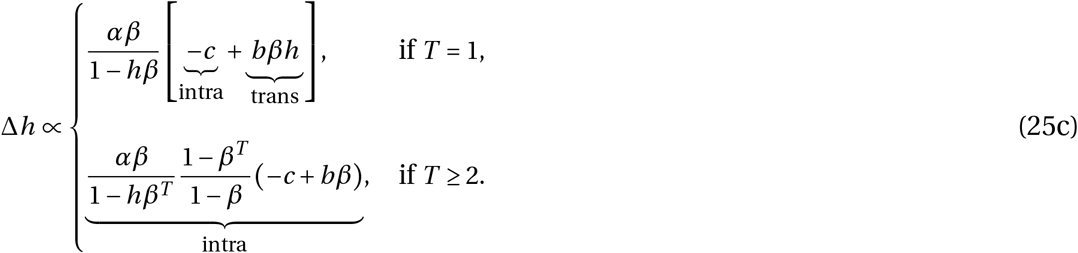

The underbraces identify the routes through which each trait affects fitness (using eq. 24).

Equations (25) provide the basis for our analyses. To clarify their implications and isolate the effect of family reputation, we first examine evolution when family reputation is absent and then compare this benchmark with scenarios in which family reputation is used.

### B.1 Evolution in the absence of family reputation

We begin with the benchmark case in which *h* = 0 is fixed, such that our model reduces to previous indirect reciprocity models (e.g. a finite-horizon version of the image-score model studied by Lehmann and Keller [3], see their Appendix S2). Selection on baseline donation and reciprocity (from eq. 25) reduces to

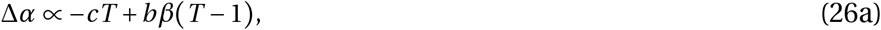

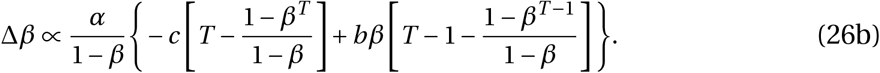

Because *h* = 0, both selection gradients contain only intragenerational effects.

Equation (26a) shows that baseline donation is favoured (i.e., Δ*α* > 0) when reciprocity in the population is greater than a threshold:

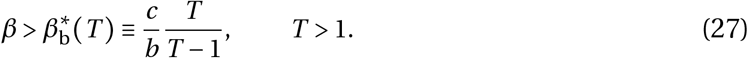

The threshold 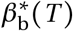 decreases with *T* and converges to *c*/*b* as *T* → ∞.

Selection on *β* (eq. 26b) is likewise positive (i.e., Δ*β* > 0) only above a critical value, denoted 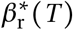 (simply referred to as *β*^*^in the main text), which is defined implicitly by

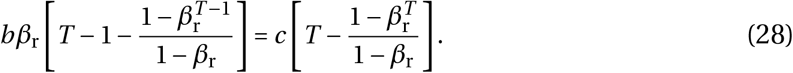

Reciprocity can evolve and be maintained only if the threshold 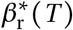is in the interior of trait space (i.e. 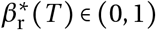). In turn, this requires that selection on reciprocity becomes positive as *β* → 1^−^. Taking this limit in eq. (26b) gives the condition

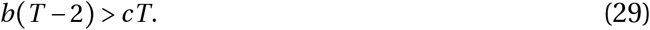

If this condition is not satisfied, reciprocity is selected against for all *β* ∈ [0, 1).

Assuming condition (29) holds true, then 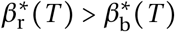 *T* holds, meaning that the condition for an increase in reciprocity is more stringent that for an increase in baseline donation. To see this, we can evaluate Δ*β* (eq. 26b) at 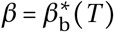, which gives a quantity with the same sign as

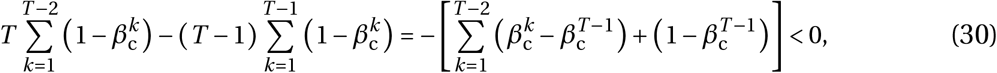

so reciprocity is selected against when 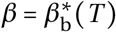. The threshold 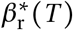 also decreases to *c*/*b* as *T* → ∞ such that

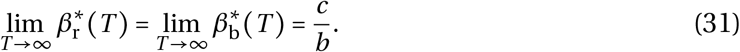

The values of this threshold in the absence of family reputation correspond to the left edge (*h* = 0) of each panel in Supplementary Fig. 1d–f. The figure shows that increasing either *T* or *b*/*c* lowers the reciprocity required for further increases in reciprocity to be favoured.

Increasing *T* therefore expands the region in which baseline donation and reciprocity reinforce one another. This is because more rounds provide more opportunities to recoup the cost of cooperation, as reputation and the resulting reciprocal donations increase over an individual’s lifetime, e.g. [3, 17].

When the resident level of reciprocity exceeds 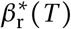, baseline donation and reciprocity reinforce one another, driving the population towards full cooperation (with *α* approaching one and *β* → 1^−^). Below this threshold, reciprocity *β* declines until it falls below 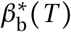, at which point baseline donation *α* also begins to decline. A lower 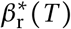 therefore enlarges the basin of attraction of cooperation and makes cooperation more robust to reductions in reciprocity in the population. Supplementary Fig. 1a–c illustrates these dynamics for *b* = 2, *c* = 1, and *T* = 4. For these parameters, condition (29) holds only at equality, so no interior threshold exists when *h* = 0 and reciprocity is selected against for every *β* < 1. Accordingly, both simulated trajectories ultimately approach the noncooperative boundary (*α* = *β* = 0).

These results align with classical models of image-score-based indirect reciprocity (e.g. [3, 18, 19]). In particular, our condition for selection to favour greater baseline donation recovers the condition for the evolution of helping under image-score-based indirect reciprocity derived by Lehmann and Keller [3] (see their Appendix S2 eq. 26). Their condition, −*C + qα*_LK_*B* > 0, is equivalent to our Δ*α* > 0 after dividing eq. (26a) by *T*, with *C* = *c, B* = *b, α*_LK_ = *β*, and *q* = (*T* − 1)/*T* .

### B.2 The effect of family reputation on the evolution of reciprocity

We now fix *h* > 0 and consider how family reputation affects the joint evolution of baseline donation *α* and reciprocity *β*.

From eq. (25a), selection favours an increase in baseline donation (Δ*α* > 0) when

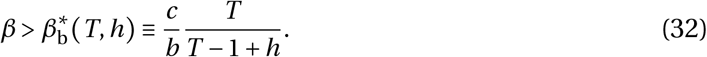

This threshold is lower than in the absence of family reputation because

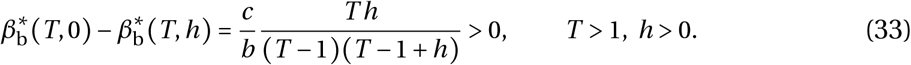

Increasing *h* lowers 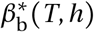 and therefore broadens the range of reciprocity values for which baseline donation increases. The reduction is due to the transgenerational benefit *bβh* in eq. (25a). An increase in baseline donation raises the mutant’s final reputation and hence the reputation inherited within its lineage (eq. 18). When *h* > 0 in the resident population, this higher inherited reputation increases the donation received by a mutant’s descendants in their first round, compensating for the absence of reputation-mediated returns during the mutant’s own first round.

Family reputation also lowers the threshold for *β* above which reciprocity increases (Δ*β* > 0), as illustrated in Supplementary Fig. 1. To show this mathematically, let *S*_*β*_ (*α, β, h, T*) denote the right-hand side of eq. (25b), so that

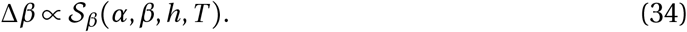

Because the proportionality factor is positive, *S* _*β*_ (*α, β, h, T*) and Δ*β* have the same sign. Let 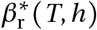 denote an interior threshold satisfying

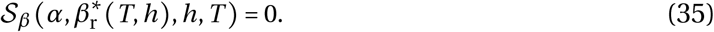

For *α* > 0, the selection gradient for reciprocity is negative when it is initially absent:

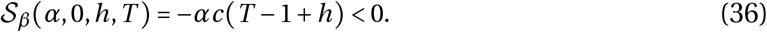

Moreover, differentiating eq. (25b) with respect to *β* gives

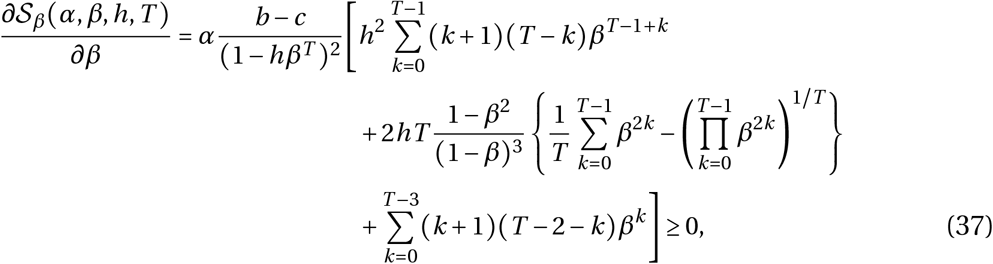

where the last sum is absent when *T <* 3. The first and last sums contain only non-negative terms. The expression in braces is also non-negative because the arithmetic mean is at least as large as the geometric mean. The derivative is therefore positive whenever an interior threshold exists. Consequently,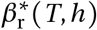 is unique: reciprocity decreases below this threshold and increases above it.

Because 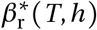 has no closed form, we determine how it changes with *h* by differentiating its defining condition in eq. (35). The implicit-function theorem gives

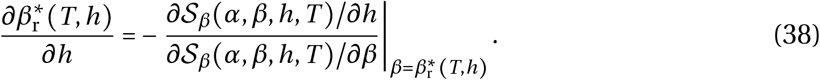

The denominator is positive by eq. (37). We therefore need to determine the sign of the numerator at the threshold. To do so, we can use

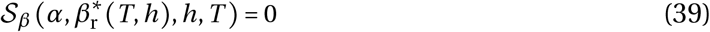

to eliminate the cost parameter *c*. Substituting the resulting expression for *c* into *∂S*_*β*_ (*α, β, h, T*) /*∂h* and simplifying, which can readily be done using a symbolic algebra program such as Mathematica, shows that

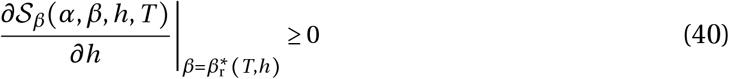

over the admissible parameter range. We omit the expanded expression because it is lengthy and provides little additional insight. Combining eqs. (38), (37), and (40) gives

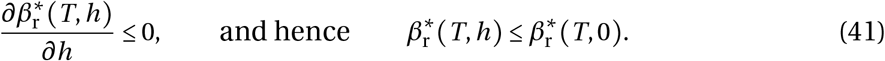

Thus, increasing reliance on family reputation lowers, or at most leaves unchanged, the amount of pre-existing reciprocity required for further increases in reciprocity to be favoured. Intuitively, stronger reciprocity raises the mutant’s final donation and therefore the reputation inherited within its lineage. When *h* > 0, this improved inherited reputation increases the donation received by descendants in their first round, adding a transgenerational return to reciprocity.

The reciprocity threshold remains at least as high as the threshold for baseline donation (i.e., 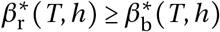. Indeed, substituting 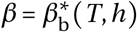 from eq. (32) into eq. (25b) gives

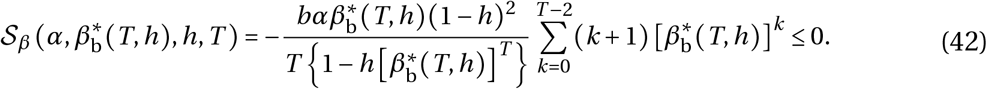

Because S_*β*_(*α, β, h, T*) increases with *β*, this implies

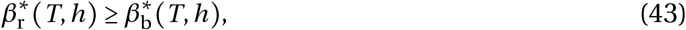

with a strict inequality when *α* > 0, *T* > 1, and *h* < 1. Hence, whenever reciprocity increases, baseline donation also increases.

Two limiting cases reveal how family reputation can substitute for repeated interactions.

First, when individuals interact only once (*T* = 1), the two thresholds coincide:

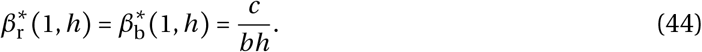

Thus, remarkably, baseline donation and reciprocity can both evolve even when individuals never interact more than once, provided that *h* > *c*/*b* and *β* > *c*/ (*bh*). Without family reputation, a donation made in a single interaction cannot generate any return. With family reputation, however, that donation determines the reputation inherited by the donor’s offspring and thereby affects the donations they receive. Family reputation therefore replaces a sequence of interactions within a lifetime with a sequence of reputation-mediated interactions across generations.

Second, when individuals rely fully on family reputation (*h* = 1), the thresholds coincide for any number of rounds:

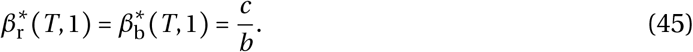

In this case, inherited reputation completely fills the informational gap in the first round: the final donation of one generation determines how its descendants are treated at the beginning of the next. Cooperation therefore faces the same threshold as under an indefinitely long interaction horizon, even when *T* is finite. The case *h* = 1 corresponds closely to the scenario studied by Albert and Rusch [20]. Using binary actions and reputations, they used simulation to show that fully inherited reputation can stabilise conditional cooperation against opportunistic strategies across both single and multiple interactions. Our model extends this insight by allowing donations, responsiveness, and reliance on family reputation to vary continuously and to coevolve.

Together, these two limiting cases clarify why the effect of family reputation depends on *T* in the baseline model. Inherited reputation is most consequential when interaction sequences are short, because the first round, in which personal reputation is not yet available, then represents a large share of an individual’s lifetime interactions. As *T* increases, this initial informational gap becomes less important, and the reduction in the thresholds produced by increasing *h* becomes smaller. Accordingly, for any fixed *h*, both thresholds converge to *c*/*b* as *T* → ∞ (eq. 31); under full reliance on family reputation, they attain this limit for every *T* . This is shown in Supplementary Fig. 1d–f: increasing *h* produces the largest reduction in 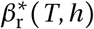 when *T* is small, its effect weakens as *T* increases, and all thresholds equal *c* /*b* when *h* = 1. This attenuation arises because the baseline model restricts the use of family reputation to the first round. In a later extension, we allow individuals to use family reputation whenever personal information is unavailable and show that its effect then persists as *T* increases.

Overall, fixing reliance on family reputation at *h* > 0 preserves the same qualitative evolutionary dynamics as when *h* = 0, but enlarges the basin of attraction of cooperation. By lowering the thresholds for both baseline donation and reciprocity, family reputation allows cooperation to evolve from less cooperative initial conditions and reduces the likelihood that a decline in reciprocity will lead the population towards non-cooperation. Supplementary Fig. 1 illustrates the resulting expansion of the cooperative basin.

### B.3 Evolution of reliance on family reputation

Having established how a fixed reliance on family reputation affects the evolution of baseline donation and reciprocity, we now allow *h* itself to evolve. For *α* > 0, eq. (25c) shows that selection favours greater reliance on family reputation (Δ*h* > 0) when

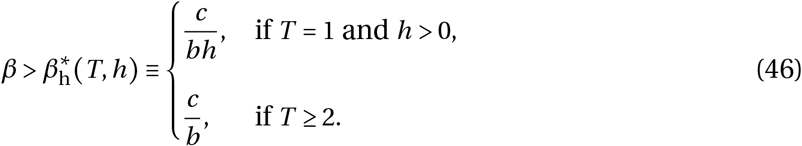

When *T =* 1, this threshold coincides with those governing the evolution of baseline donation and reciprocity:

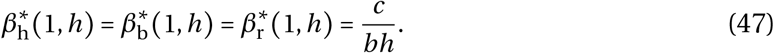

Thus, all three traits can increase even though each individual interacts only once. The threshold can be reached when *h* > *c*/*b*; in which case if *β* > *c*/(*bh*), the evolution of the three traits becomes mutually reinforcing (see eq. 25). Greater baseline donation creates a better reputation to transmit, greater reciprocity increases the benefit that descendants receive from inheriting this reputation, and greater reliance on family reputation strengthens this intergenerational return. This positive feedback drives baseline donation, reciprocity, and reliance on family reputation towards the fully cooperative state. This shows that a pre-existing tendency to use family reputation can allow reciprocity to evolve in a one-shot setting, with stronger reliance making cooperation easier to establish and maintain.

When *T* ≥ 2, eq. (46) shows that reliance on family reputation can evolve from *h* = 0, provided that *α* > 0 and resident reciprocity exceeds *c*/*b*. Higher *h* increases the donor’s first-round donation, thereby improving its personal reputation and generating a reciprocal benefit in the following round. Unlike when *T* = 1, reliance on family reputation therefore need not already be established to evolve. Whenever the interior reciprocity threshold 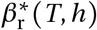 exists, the three thresholds satisfy

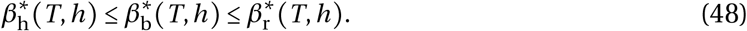

The first inequality follows from *T / (T* − 1 + *h)* ≥ 1 (eq. 32), whereas the second was established in eq. (43). Consequently, whenever selection favours greater reciprocity, it also favours greater baseline donation and greater reliance on family reputation. The three traits therefore reinforce one another throughout the cooperative basin: increasing *h* further lowers the thresholds for baseline donation and reciprocity, expanding the cooperative basin and making cooperation more robust to declines in reciprocity.

These results show why reliance on family reputation is generally favoured in cooperative populations. Before an individual has interacted, family reputation provides information about its likely behaviour. Because cooperative traits are transmitted between generations, parental reputation is predictive of the behaviour expected from offspring. Placing greater weight on this information allows individuals to condition their first interaction on an informative signal, rather than on an initial reputation that is unrelated to the partner’s likely behaviour.

The route through which this information produces a fitness benefit depends on the number of rounds. When *T* = 1, a mutant with a higher *h*_m_ makes a larger donation in its only interaction, provided that *α* > 0 and *β* > 0 (eq. (16)). This donation is also its final donation and therefore determines the reputation inherited within its lineage (eq. (18)). The mutant pays the cost of donating more but cannot receive a return during its own lifetime. Its descendants may nevertheless benefit from inheriting its improved reputation. Selection on *h* therefore combines an intragenerational cost, −*c*, with a transgenerational benefit, *bβh* (eq. 25c).

When *T* ≥ 2, the mutant’s final donation no longer depends on *h*_m_, because it is based on the resident’s personal rather than inherited reputation. Changes in *h*_m_ therefore do not affect the reputation transmitted by the mutant, and selection on *h* has no transgenerational component (eq. 25c). Instead, the mutant’s larger first-round donation improves its personal reputation and increases the donation it receives in the following round. Greater reliance on family reputation is consequently favoured whenever the resulting reciprocal benefit, *bβ*, exceeds the cost *c*.

The evolution of *h* therefore generates a positive feedback with cooperation. As cooperation and reciprocity increase, family reputation becomes more valuable and selection favours placing greater weight on it. In turn, increasing *h* lowers both 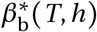 and 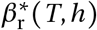 (eqs. (33) and (41)). Baseline donation and reciprocity can then continue to increase at progressively lower levels of resident reciprocity. A population with a higher *h* can therefore withstand a larger reduction in *β* before crossing into the region where reciprocity declines. The comparisons across fixed values of *h* in Supplementary Fig. 1 illustrate how this feedback enlarges the basin of attraction of cooperation and makes the cooperative state more robust.

These results assume that using family reputation entails no cognitive cost ([21]) and that family reputation is known perfectly. A cognitive cost or imperfect information would of course reduce the value of family reputation, weaken selection for greater *h*, and narrow the conditions under which it evolves.

## C Individual-based simulations

We use individual-based simulations to test whether the analytical predictions hold in finite populations subject to mutation and genetic drift, and to examine extensions of the baseline model. These extensions include errors in donation and reputation assessment, incomplete observation of behaviour, imperfect knowledge of family reputation, and reputations shared among family members. We first describe the baseline simulation and then introduce each extension in turn. All simulation code is publicly available from the Open Science Framework repository https://osf.io/frpyh/.

### C.1 Baseline

#### C1.1 Model and algorithm

The baseline simulation follows the analytical model in Section A, but considers a finite population with stochastic reproduction and mutation. The population consists of *N =* 1000 individuals with non-overlapping generations. Supplementary Table S1 summarizes the simulation parameters.

Each individual *i* ∈ {1,…, *N* } is characterized by the phenotype

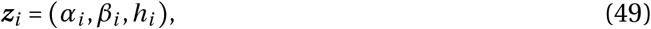

where *α*_*i*_ is baseline donation, *β*_*i*_ is reciprocity, and *h*_*i*_ is reliance on family reputation. Individual *i* also carries an inherited family reputation *ρ*_F,*i*_, equal to its parent’s final donation in the preceding generation (eq. 4). At birth, personal reputation is initialized at the default value *ρ*_ini_ = 0 (e.g., as in [6]), whereas family reputation remains available until a personal reputation has been established.

Unless stated otherwise, simulations are initialised with a monomorphic cooperative population in which

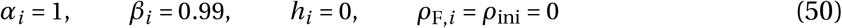

for every individual *i* . This initialization allows us to test whether cooperation remains evolutionarily stable and whether reliance on family reputation can evolve from zero.

Each generation proceeds through the following steps.

##### 1. Social interactions

Individuals participate in *T* rounds of the donation game. At the beginning of each round, the population is randomly shuffled and divided into *N*/2 pairs. Each individual therefore interacts exactly once per round, with partners independently rematched between rounds.

Consider individual *i* paired with individual *j* in round *t* ∈ {1,…, *T}*. The reputation of *j* used by *i* is

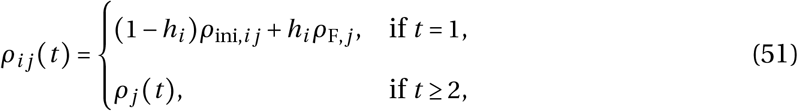

where *ρ*_*j*_ *t* is *j* ‘s public personal reputation at the beginning of round *t*, and *ρ*_ini,*ij*_ is the initial reputation that *i* assigns *j* . In the baseline model, *ρ*_ini,*ij =*_ *ρ*_ini_ = 0, but we later relax this assumption. The intended donation from *i* to *j* is then

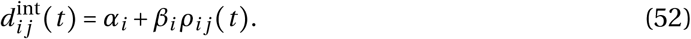

The simulation allows for errors in donation execution. Independently for every player and interaction, an execution error occurs with probability _e_. The realised donation is

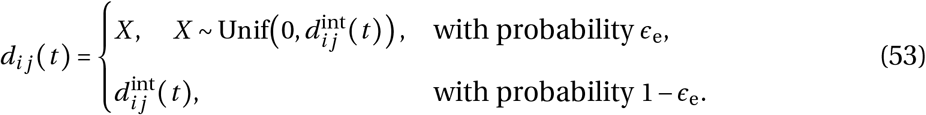

Execution errors can therefore reduce an intended donation but cannot increase it, consistent with the standard assumption that intended cooperation may fail [22].

The realised donations update the accumulated payoffs of the two players according to

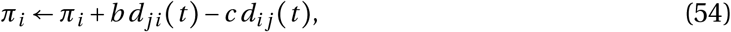

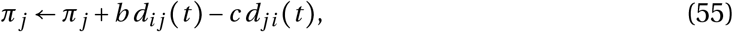

with *π*_*i*_ = *π*_*j*_ = 0 at the beginning of each generation.

##### 2. Reputation updating

After each interaction, the public personal reputation of each player is updated from its realised donation. With probability 1 − ϵ_a_, the donation is assessed correctly. With probability _a_, an assessment error occurs and the new reputation is drawn uniformly from the feasible reputation interval. Thus,

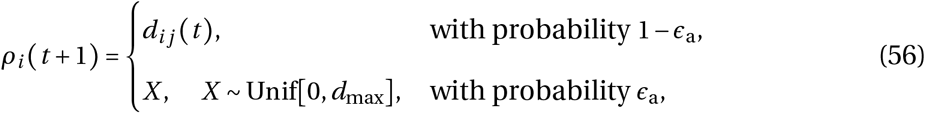

where

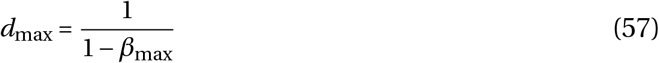

is the maximum feasible donation in our simulation (given by the limit of Eq. 11 when *T* → ∞), and *β*_max_ = 0.99. A single public assessment is made for each donation and is shared by the entire population. Family reputations remain unchanged during the *T* interaction rounds.

#### 3. Reproduction

After the final round, the raw lifetime payoff of individual *i* is *π*_*i*_ . Fecundity is then *f*_*i*_ = 1 + *π*_*i*_ . To ensure non-negative fecundities, these fecundities are shifted by a common constant whenever at least one individual has a negative fecundity: 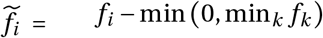 . The common fecundity shift preserves the ordering of individuals by fecundity, while keeping fecundities non-negative. The case when all individuals have zero fecundity never occurs since *b* ≥ *c* in our simulation.

The next generation is formed by independently sampling *N* parents with replacement. Individual *i* is chosen as the parent of each offspring with probability

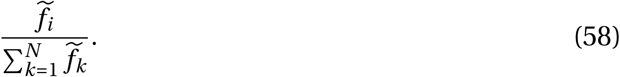

Each offspring inherits all three traits from its sampled parent. It also inherits the parent’s final donation as its family reputation, so all offspring of the same parent receive the same *ρ*_F_. Adults then die and the offspring become the adults of the next generation.

##### 4. Mutation

For each offspring, the three traits mutate independently. A given trait mutates with probability *μ* = 0.001. Conditional on mutation, a normally distributed perturbation is added to the inherited value:

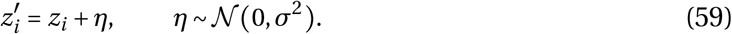

Baseline donation and reliance on family reputation are clipped to remain in *α*_*i*_, *h*_*i*_ ∈ [0, 1], whereas reciprocity is clipped to remain to *β*_*i*_ ∈ [0, 0.99]. Restricting *β*_*i*_ below one prevents donations from increasing without bound. When a trait is held fixed in a particular analysis, mutation of that trait is disabled. The mutational standard deviation *σ* is reported in the corresponding figure legend; the main-text simulations use *σ* = 0.1.

Each simulation runs for 10^6^ generations. Population mean trait values are recorded every 100 generations, and their long-term values are calculated by averaging the recorded values over the final third of the simulation. Parameter values that differ among analyses are reported in the corresponding figure legends.

#### C.1.2 Results

We first asked whether the simulations recover the analytical predictions in the absence of family reputation. When *h* = 0 is fixed, an initially cooperative population remains cooperative only when *T* and *b* are sufficiently large (Supplementary Fig. 2, top row), consistent with the condition in eq. (29). Increasing either the number of rounds or the benefit of donation expands the parameter region in which cooperation is maintained.

We next examined how a fixed reliance on family reputation changes the basin of attraction of cooperation. Supplementary Fig. 1a–c compares simulated trajectories with the evolutionary directions predicted from the analytical selection gradients (eq. 25). The streamlines represent the predicted trajectories obtained from Lande’s equation [23]:

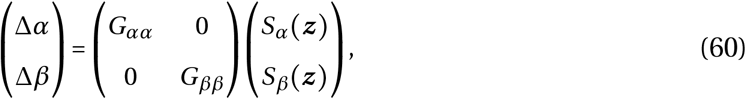

assuming independent genetic variation in *α* and *β*. We set *G*_*αα*_= *G*_*ββ*_, so the direction of each streamline is determined by the selection-gradient vector. The points in Supplementary Fig. 1a–c show population means from individual-based simulations.

For the parameters used in Supplementary Fig. 1a–c, the critical reciprocity threshold is 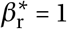 when *h*=0 and therefore lies outside the simulated trait range, *β*≤ *β*_max_ =0.99 (Supplementary Fig. 1a). Reciprocity consequently declines from every feasible initial value. Once reciprocity falls below the threshold for baseline donation (*β*_b_ =2/3), *α* also declines and the population evolves towards defection. When *h* =0.5, the reciprocity threshold falls to 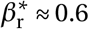: populations starting above this threshold evolve towards cooperation, whereas those starting below it evolve towards defection (Supplementary Fig. 1b). Under full reliance on family reputation (*h*=1), the threshold reaches its minimum, 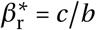 (eq. 45), further enlarging the cooperative basin (Supplementary Fig. 1c). The simulated trajectories closely follow these predicted directions.

We then allowed *h* to evolve from an initial value of zero. Reliance on family reputation evolves across most of the parameter space explored (Supplementary Fig. 2, bottom row), with the main exception occurring when the net benefit of cooperation are very small (*b* < 1.05). Comparing simulations in which *h* is fixed at zero with those in which it can evolve shows that the evolution of *h* expands the parameter region in which cooperation is maintained (compare top and bottom rows of Supplementary Fig. 2). This effect is strongest when interactions are few and *b* only moderately exceeds *c*, where personal reputation alone provides insufficient returns to sustain cooperation. Where cooperation is already maintained without family reputation, the evolution of *h* nevertheless increases average donations. By carrying informative reputations across generations, increasing *h* reinforces selection on baseline donation and reciprocity rather than requiring individuals to begin each generation from an uninformative initial reputation.

In Figure Fig. 3a of the main text, we examined the joint evolutionary dynamics of reciprocity (*β*) and reliance on family reputation (*h*), while keeping the baseline donation level fixed (*α* =0.1). This figure again compares simulated trajectories with the evolutionary directions predicted from the analytical model, with streamlines represent the predicted trajectories obtained from Lande’s equation [23]:

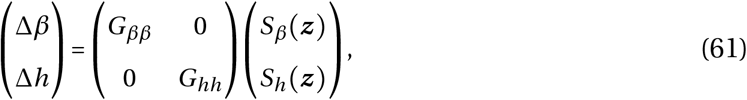

assuming independent genetic variation in *β* and *h*. We set *G*_*ββ*_= *G*_*hh*_, so the direction of each streamline is determined by the selection-gradient vector. The points in Fig. 3a show population means from individual-based simulations.

We tested whether these conclusions remain robust to errors in donation execution and reputation assessment. When individuals interact few times (*T*=5), family reputation substantially expands the range of error rates under which cooperation persists: an initially cooperative population remains cooperative when both ϵ_e_ and ϵ_a_ are approximately 0.35 (Supplementary Fig. 3). When individuals interact more often (*T* = 20), cooperation remains stable over a similar range of errors with or without family reputation (Supplementary Fig. 4). Nevertheless, family reputation evolves and continues to increase average donations. Thus, family reputation is most effective at preventing the collapse of cooperation when individuals interact only a few times, but it also increases average donations when cooperation would persist without it.

Finally, we also relaxed the assumption that individuals assign a default reputation of zero. We now assume that the initial reputation is random [8]. To do so, at the beginning of each generation, the initial default reputation *ρ*_ini,*ij*_ that individual *i* assigns to partner *j* is drawn uniformly between 0 and the maximum feasible donation (*d*_max_). Thus,

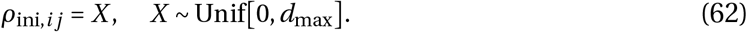

At the beginning of a simulation, family reputations were similarly initialized with a random *ρ*_ini_.

Supplementary Fig. 5 confirms that reliance on family reputation *h* still evolves in most of the parameter space in this scenario. However, unlike before, allowing *h* to evolve does not expand the region where cooperation is stable (Supplementary Fig. 5, compare top and bottom rows). Individuals now start with an expected initial reputation that is positive and large (i.e., *d*_max_/2), which weakens the advantage of using positive family information compared to initial reputations of zero. Another difference from the baseline model is that *h* evolves through drift or is even counter-selected with a low number of rounds (*T*<5; Supplementary Fig. 5). The reason is that, when *h* is initially low or absent, reputations need several rounds to decorrelate from the random initial draw (Eq. 62) and start reflecting genotypes. Nevertheless, when it evolves, reliance on family reputation again increases donations (Supplementary Fig. 5).

### C.2 Imperfect reputation knowledge

#### C.2.1 Model

We next relax the assumption that every interaction is publicly observed. Each interaction is now independently public with probability *q* and private with probability 1−*q*. Following a public interaction, the personal reputations of both players are updated from their realised donations, subject to the assessment errors described above (section C.1.1). Following a private interaction, neither reputation is updated. An individual’s personal reputation is therefore unavailable until at least one of their interactions has been publicly observed; thereafter, it records their most recent publicly assessed donation.

When individual *i* interacts with individual *j* in round *t*, the reputation used by *i* is

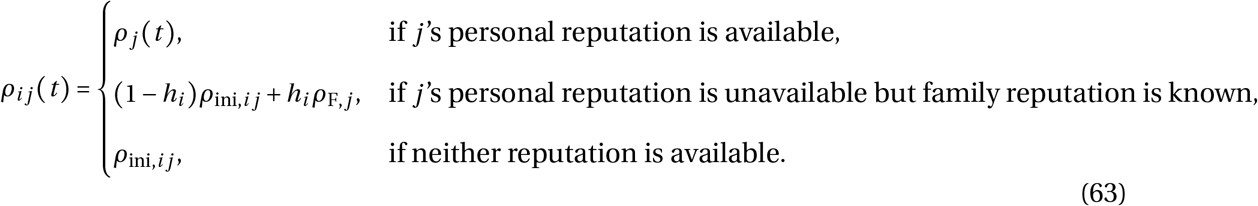

Here, *ρ*_*j*_ *t* is *j*‘s most recent publicly assessed donation and *ρ*_ini,*ij*_ *ρ*_ini_ 0 is the default reputation. Whenever *j*‘s personal reputation is unavailable, *i* knows *j*‘s family reputation independently with probability *s*. Family reputation remains fixed within a generation.

By round *t*, individual *j* has participated in *t*−1 previous interactions. Their personal reputation remains unavailable only if all these interactions were private, which occurs with probability (1–*q)*^*t*−1^. The expected proportion of rounds in which a partner’s personal reputation is unavailable is therefore

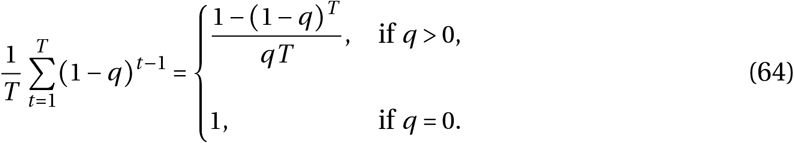

This proportion decreases with both observability *q* and the number of rounds *T*. Thus, family reputation is available as an alternative source of information most often when public observations are rare or individuals interact only a few times.

#### C.2.2 Results

We first asked whether reliance on family reputation evolves when personal behaviour is imperfectly observed. Across most combinations of *q* and *s, h* increases from its initial value of zero and cooperation is maintained (Supplementary Fig. 8). When family reputation is sufficiently often known, reliance on family reputation compensates for lower observability, because personal reputation remains unavailable for more interactions (eq. (64); compare top and bottom rows of Supplementary Fig. 8). The main exception occurs when both public observations and knowledge of family reputation are rare (Supplementary Fig. 8, bottom row). Under these conditions, neither source provides sufficient information about partners, and cooperation collapses.

Where reliance on family reputation evolves, average donations remain high even when few interactions are publicly observed (Supplementary Fig. 8, compare top and bottom rows). Family reputation compensates for the missing personal information by carrying reputational information across generations. Because an individual’s family reputation equals its parent’s final donation, the increase in donations generated by reciprocity can continue from one generation to the next. When *h* approaches one, reputations can therefore approach the equilibrium *ρ*^∗^=*α*/(1−*β*) even at low observability (eq. (11); Supplementary Fig. 9). By contrast, when *h* remains low, individuals begin each generation near the default reputation and must build their personal reputation through publicly observed interactions. When both *q* and *T* are small, too few such observations occur for reputations to approach equilibrium within a lifetime (Supplementary Fig. 9).

Thus, provided that family reputation is sufficiently often known, its evolution maintains cooperation and increases donations when personal observations are scarce.

### C.3 Single, shared family reputation between family members

#### C.3.1 Model

We next allow the behaviour of one individual to affect how all its siblings are evaluated. Following reproduction, all offspring produced by the same parent are assigned to a common sibling group. We denote the group containing individual *i* by *g*(*i*). At the beginning of the generation, all members of a sibling group share the family reputation inherited from their parent. At the start of a simulation, each individual is alone in its sibling group.

During the generation, the shared family reputation is updated whenever the donation of any sibling is publicly observed. Specifically, the publicly assessed donation made by individual *i* becomes the family reputation of every member of *g* (*i*). The personal reputation of *i* is updated at the same time, as in the preceding model. Private interactions update neither personal nor family reputation. Thus, at any point in the generation, *ρ*_F,*g* (*i*_) (*t*) records the most recent publicly assessed donation made by any member of *i* ‘s sibling group. Consequently, a more cooperative donation by one sibling improves the reputation of all siblings, whereas a less cooperative donation harms it.

To isolate the consequences of sharing a family reputation, we consider a deliberately simple decision rule in which individuals rely exclusively on either family or personal reputation throughout life. At the beginning of its life, each individual *i* adopts one of two information sources:

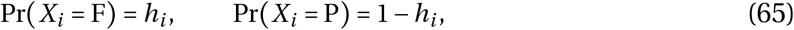

where *X*_*i*_= F denotes exclusive reliance on family reputation and *X*_*i*_= P denotes exclusive reliance on personal reputation. This information source remains fixed throughout the individual’s lifetime. Thus, in this extension, *h*_*i*_ is the probability of relying exclusively on family reputation rather than a continuous weight assigned to it.

As in the preceding model, family reputation is known independently with probability *s* in each interaction. The reputation that individual *i* assigns to partner *j* is therefore

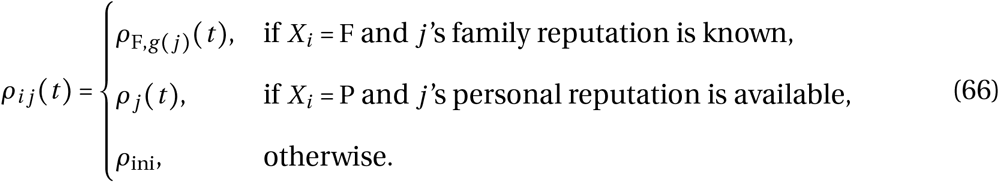

An individual relying on family reputation therefore ignores the partner’s personal reputation even when it is available, whereas an individual relying on personal reputation does not use family information.

This reputation system is analogous to the out-group homogeneity bias, whereby members of another social group are perceived as more similar than they are [24], which might be the basis for stereotypes [21, 22]. Here, groups are defined by common parentage rather than by broader social categories. The model represents a strong form of this effect because individuals relying on family reputation evaluate all siblings identically, despite any differences in their behaviour or traits.

#### C.3.2 Results

We asked whether reliance on family reputation can evolve when family members share a common reputation and individuals use either family or personal information throughout life. It can, although under a narrower range of conditions than in the preceding model (compare Fig. 4a and b). As before, fewer rounds and lower observability favour reliance on family reputation. Under low observability, however, *h* increases only when family reputation is sufficiently often known (high *s*; Fig. 4b).

This stronger dependence on *s* follows from the exclusive use of one information source.

An individual relying on family reputation ignores personal reputation throughout life. If the partner’s family reputation is unknown, the individual must therefore use the default reputation even when personal information is available. When *s* is low, this loss of personal information outweighs the benefit of using family reputation, and *h* can be selected against even in cooperative populations. By contrast, in the preceding model, unavailable family information matters only until the partner’s behaviour is first observed.

When family reputation is sufficiently often known, *h* evolves and average donations increase substantially (Fig. 4b). Each publicly observed donation then updates the reputation shared by the donor’s entire sibling group, allowing the behaviour of one family member to affect how all siblings are treated. A shared family reputation can therefore compensate for limited observations of personal behaviour and support high levels of cooperation, but its evolution requires family information to be widely available.

### C.4 Extended classic pure-strategies indirect reciprocity game

#### C.4.1 Model

We finally examine whether our results extend to the discrete strategies commonly considered in models of indirect reciprocity [6, 8, 25]. The population structure, reproduction, and inheritance follow the baseline simulation described above (section C.1.1). Here, we describe the changes to the interaction and mutation rules.

Donations are binary, and each generation contains *T* rounds. As before, each individual participates in exactly *T* interactions, and each individual in a pair can decide to cooperate or not. Cooperation incurs a cost *c* and provides the recipient with a benefit *b*, whereas defection has no effect on either payoff. Thus, if *d*_*ij*_ (*t*)∈{0, 1} denotes the action of player *i* towards player *j*, payoffs are updated according to

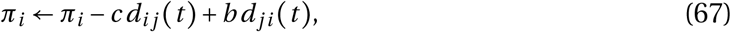

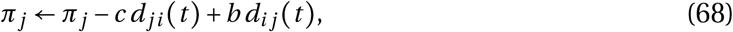

where *d*_*ij*_ (*t*) = 1 denotes cooperation and *d*_*ij*_ (*t*) = 0 denotes defection.

Reputations are also binary and follow image scoring with public assessment [7]. Following a publicly observed interaction, a donor is assigned a GOOD reputation after cooperating and a BAD reputation after defecting. A private interaction does not update reputation. Individuals whose behaviour has not yet been observed are assigned a random default reputation *ρ*_ini_ uniformly drawn from {BAD, GOOD} [8], unless stated otherwise. Offspring inherit their parent’s final binary reputation as their family reputation. Family reputation was also randomly initialized at the beginning of a simulation.

Individuals express one of four strategies: (i) ALLD always defects; (ii) ALLC always cooperates; (iii) Disc cooperates with recipients having a GOOD personal reputation and defects against recipients having a BAD personal reputation. If the recipient has not yet been observed, Disc uses the default reputation; (iv) DiscH follows the same conditional rule as Disc but, when the recipient has not yet been observed, uses the recipient’s inherited family reputation. This family reputation is known with probability *s*; otherwise, DiscH uses the default reputation.

When errors are included, intended cooperation is incorrectly implemented as defection with probability _e_, whereas intended defection remains unchanged. Following an observed interaction, the donor’s assigned reputation is reversed with probability _a_. We consider positive error rates by default, as is often the case in indirect reciprocity models [8, 25].

Strategies are transmitted from parent to offspring. With probability *µ*, an offspring mutates to another strategy, chosen uniformly from the other strategies available in that simulation. We compare two conditions. In the three-strategy condition, only ALLD, ALLC, and Disc are available, recovering the classical model [6]. In the four-strategy condition, DiscH is also available. Comparing these conditions tests whether discriminators using family reputation evolve and how their presence affects cooperation.

All simulations are initialised with a population composed entirely of ALLD individuals. Each run lasts 10^6^ generations. Strategy frequencies and the proportion of donor decisions resulting in cooperation are recorded every 100 generations, and long-term averages are calculated over the final third of each run.

#### C.4.2 Results

We first consider the three-strategy condition in which DiscH cannot evolve. Within the parameter range explored, substantial cooperation evolves only when the benefit of cooperation is high (*b* > 4; Supplementary Fig. 6, top row). Even then, cooperation does not settle at a stable level (Fig. 3c). Instead, strategy frequencies undergo the cycles characteristic of image-scoring models [18, 26]. Discriminators spread and suppress ALLD by withholding cooperation from individuals with a BAD reputation. Once defectors become rare, ALLC and Disc behave similarly and ALLC can increase neutrally through drift. The resulting abundance of unconditional cooperators then allows ALLD to spread, restarting the cycle. Consequently, cooperation repeatedly rises and collapses (Fig. 3c), and the three strategies have broadly similar long-term average frequencies where cooperation evolves (Supplementary Fig. 6, top row).

Allowing DiscH to evolve increases both the prevalence and stability of cooperation (compare Supplementary Fig. 6 top and bottom rows). Cooperation is maintained at lower benefits (*b*>2.5) and with fewer rounds than in the three-strategy condition (Supplementary Fig. 6, bottom row). The effect is strongest when *T* is small. Remarkably, cooperation can evolve even when individuals interact only once (*T*=1), provided that the benefit is sufficiently large. This reproduces the central result of our continuous-trait model: inherited reputation can generate reputational returns across generations when such returns cannot arise through repeated interactions within a lifetime.

Family reputation particularly benefits ALLC. Unconditional cooperators are the strategy most likely to end life with a GOOD reputation and transmit it to their offspring. By contrast, a discriminator may end with a BAD reputation after refusing to help a recipient with a BAD reputation, because image scoring treats every defection negatively. By using inherited reputation, DiscH therefore preferentially helps the descendants of unconditional cooperators before their own behaviour has been observed. Conversely, the descendants of defectors are much more likely to inherit a BAD reputation and therefore receive less help. This pushes the population away from ALLD and towards cooperative strategies.

Family reputation also dampens the cycles observed in its absence. When DiscH cannot evolve, cooperation repeatedly moves between values close to zero and one (Fig. 3c). When DiscH is present, ALLC, Disc, and DiscH form a persistent mixture, ALLD remains rare, and cooperation stays high apart from brief declines (Fig. 3c). Inherited reputation limits the ability of defectors to exploit cooperative populations because the reputational consequences of defection are transmitted to their descendants.

These effects remain when unknown individuals are initially assigned a GOOD reputation and when interactions are imperfectly observed (Supplementary Fig. 6, Supplementary Fig. 7, and Supplementary Fig. 10). Assigning a GOOD reputation to unknown partners corresponds to the positive “prejudice of discriminators” studied by Nowak and Sigmund [26]. Thus, the stabilising effect of family reputation does not depend on continuous traits or mutations of small effect: it also arises in a classical discrete-strategy model and is strongest when interactions, benefits, or observations are limited.

## Supplementary Figures

**Supplementary Figure 1.**
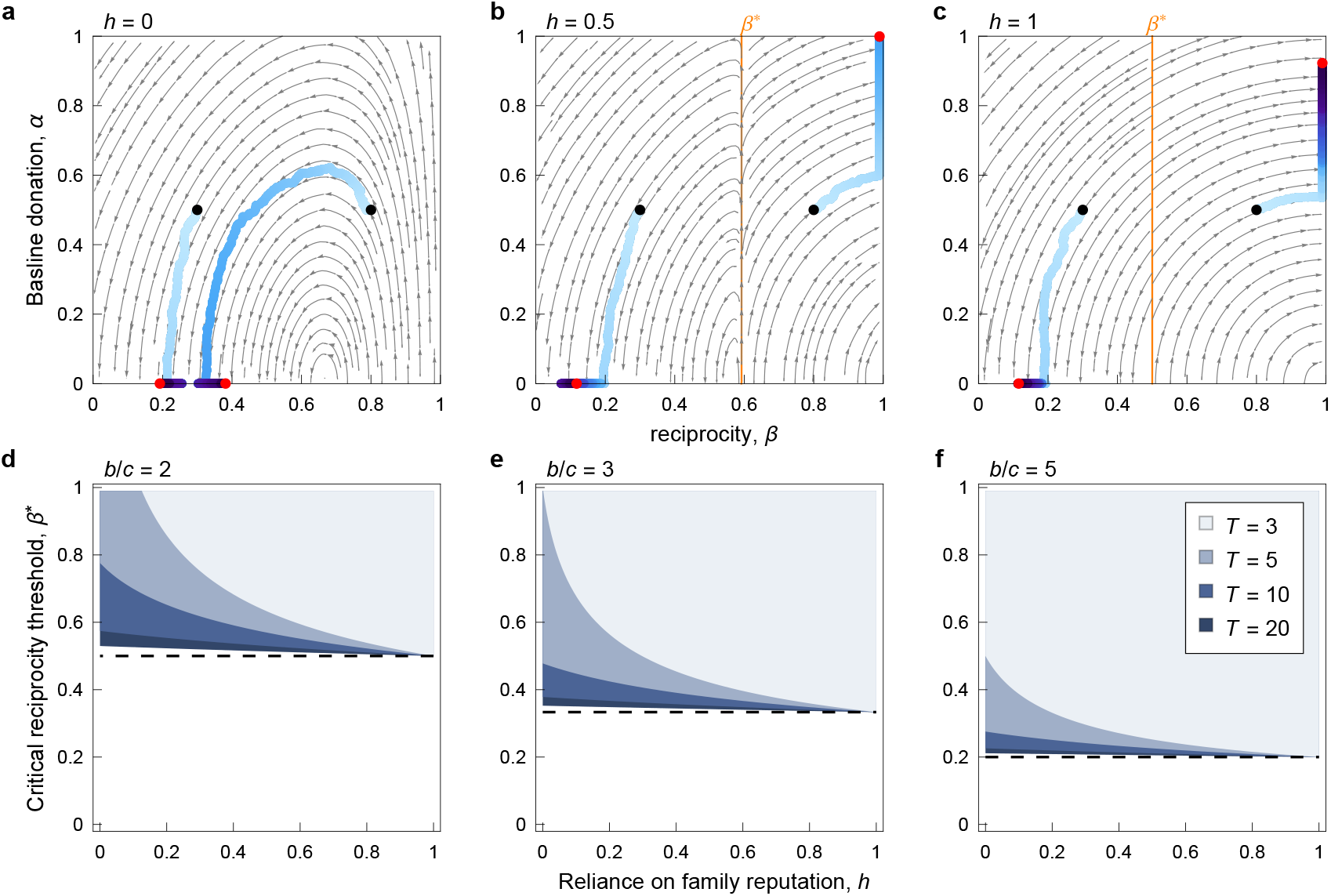
Reliance on family reputation favours reciprocity. **a–c**, Joint evolutionary dynamics of baseline donations *α* and reciprocity *β* (eq. 60), when reliance on family reputation (*h*) is fixed in the population. Blue dots show the observed trajectories in simulation runs (darker blues indicate later generations; see Section C.1 for simulation details). Black and red dots indicate the initial trait values and final ones after 3×10^6^ generations, respectively. The panels show different values of *h*. Greater reliance on family reputation stabilises cooperation, by lowering the critical reciprocity threshold *β*^∗^ above which further increases in reciprocity are favoured (orange line). **d–f**, Critical reciprocity threshold *β*^∗^above which further increases in reciprocity are favoured, as a function of reliance on family reputation (*h*), for different benefit-to-cost ratios (*b*/*c*) and number of rounds (*T*). Greater reliance on family reputation in the population reduces *β*^∗^, expanding the basin of attraction of cooperation. The effect of *h* on *β*^∗^ decreases as the number of rounds increases. The dashed line represents *β*^∗^in the limit of many interactions (*c*/*b*). The threshold *β*^∗^ is found by solving the zeroes of eq. 25b for *β*. Parameters: all panels, *c* = 1; panels **a–c**, *N*=10^4^, *b* = 2, *T* = 4, *q* = *s* = 1, *ρ*_ini_ = 0, ϵ_a_ = ϵ _e_ = 0, *µ* = *σ* = 10^−3^.

**Supplementary Figure 2.**
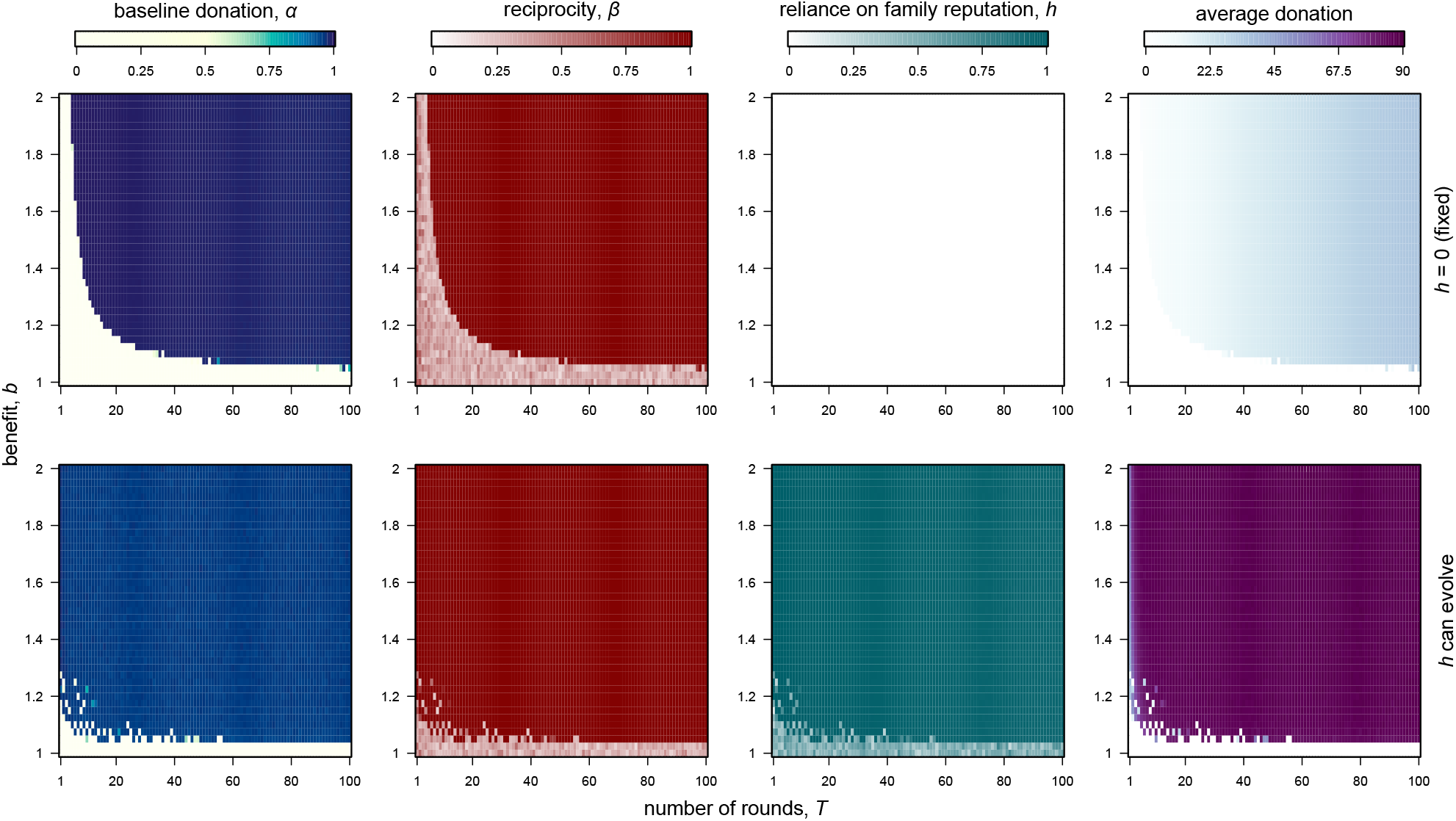
The evolution of reliance on family reputation increases the stability of cooperation. Long-term trait averages across different numbers of rounds (*T*) and benefits (*b*). Reliance on family reputation *h* was either fixed at zero or allowed to evolve (top and bottom row, respectively; see Section C.1 for simulation details.). Reliance on family reputation stabilises cooperation, especially when the number of rounds (*T*) and benefits (*b*) are small, and also improves donations. Populations were initialised in the fully cooperative state without reputation inheritance (α=1, *β* = 0.99, *h* = 0). Parameters: *N* = 10^3^, *c* = 1, *q* = *s* = 1, ϵ_a_ = ϵ_e_ = 0, *ρ*_ini_ = 0, *µ* = 10^−3^, *σ* = 0.1.

**Supplementary Figure 3.**
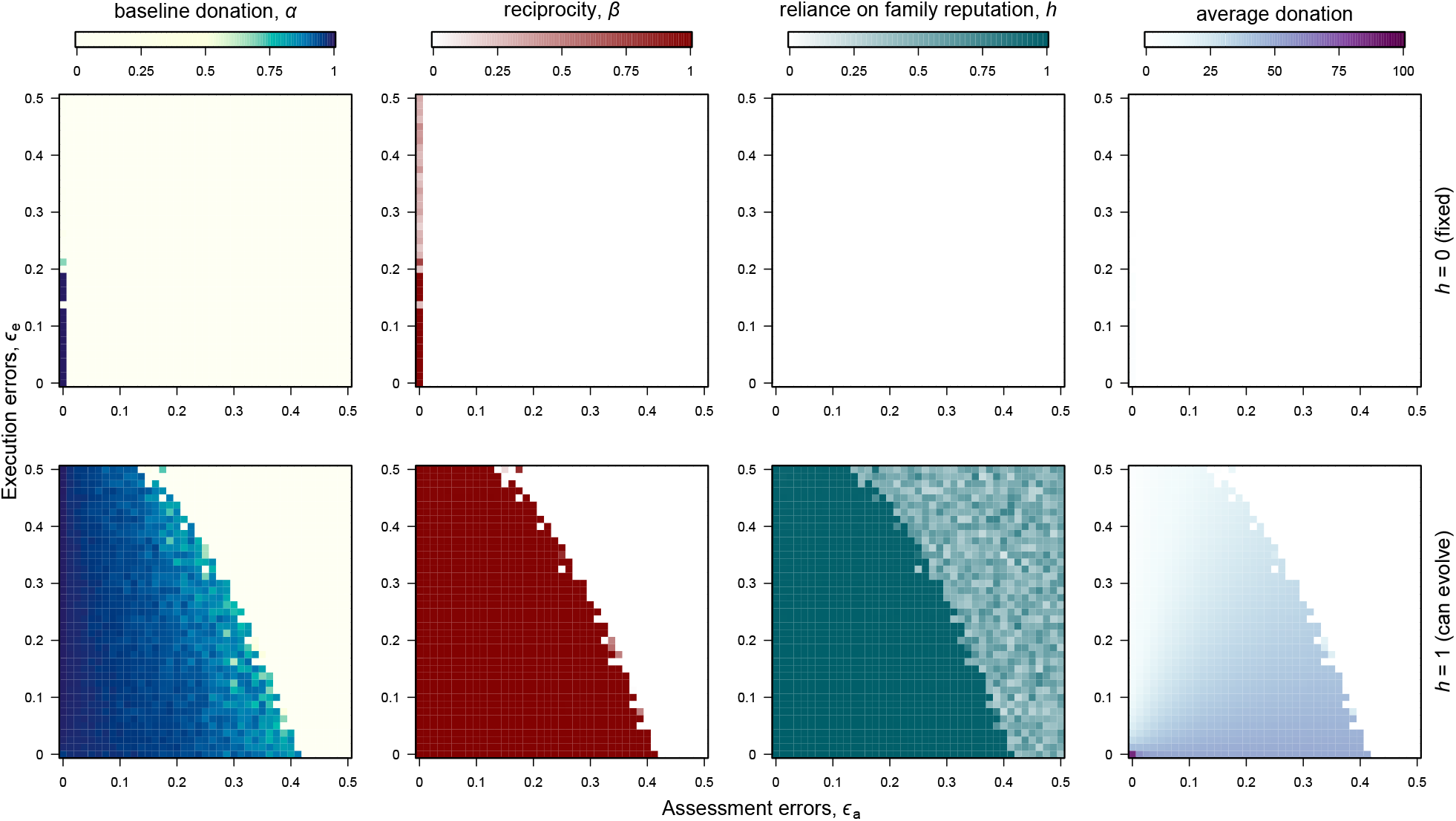
Reputation inheritance stabilises cooperation in the presence of errors. Long-term trait averages across different assessment and execution errors (ϵ_a_ and ϵ_e_, respectively). In the top row, reliance on family reputation could not evolve (*h*=0). In the bottom row, reliance on family reputation started at *h*=1 and was allowed to evolve (see Section C.1 for simulation details). When populations fully rely on family reputation (*h*=, bottom row), cooperation is maintained under higher rates of errors in both reputation assessment and action execution. Parameters: *N* = 10^3^, *c* = 1, *b* = 2, *T* = 5, *ρ*_ini_ = 0, *µ* = 10^−3^, *σ* = 0.1.

**Supplementary Figure 4.**
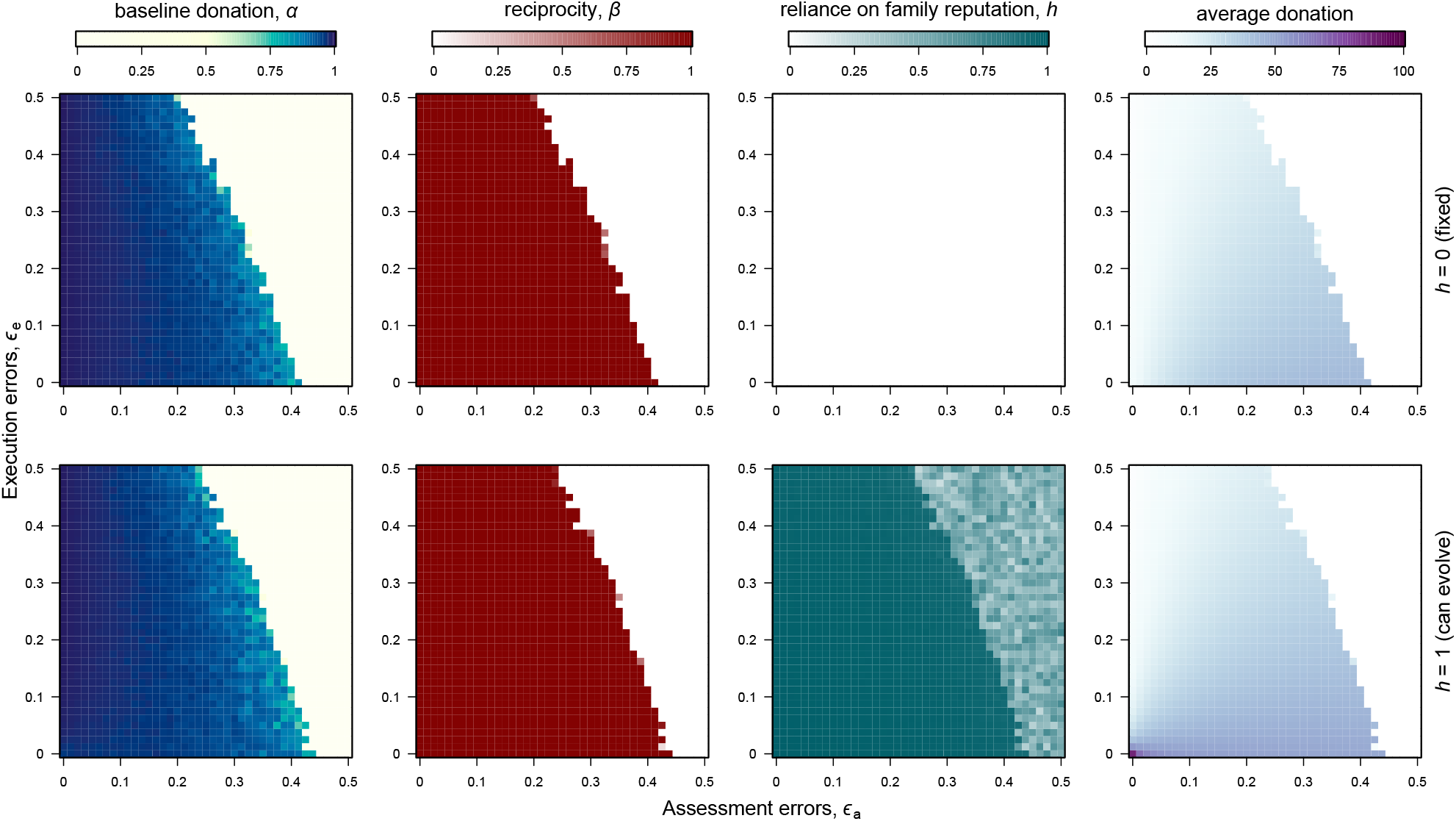
Reputation inheritance increases donations in the presence of errors. Long-term trait averages across different assessment and execution errors (ϵ_a_ andϵ_e_, respectively; see Section C.1 for simulation details). In the top row, reliance on family reputation could not evolve (*h* = 0). In the bottom row, reliance on family reputation started at *h* = 1 and was allowed to evolve. The number of rounds (*T*=20) is sufficient to maintain reciprocity without family reputation, and the presence of reliance on family reputation (bottom row) does not expand the conditions where cooperation is maintained, but still leads to higher donations. Parameters: *N* = 10^3^, *c* = 1, *b* = 2, *T* = 20, *ρ*_ini_ = 0, *µ* = 10^−3^, *σ* = 0.1.

**Supplementary Figure 5.**
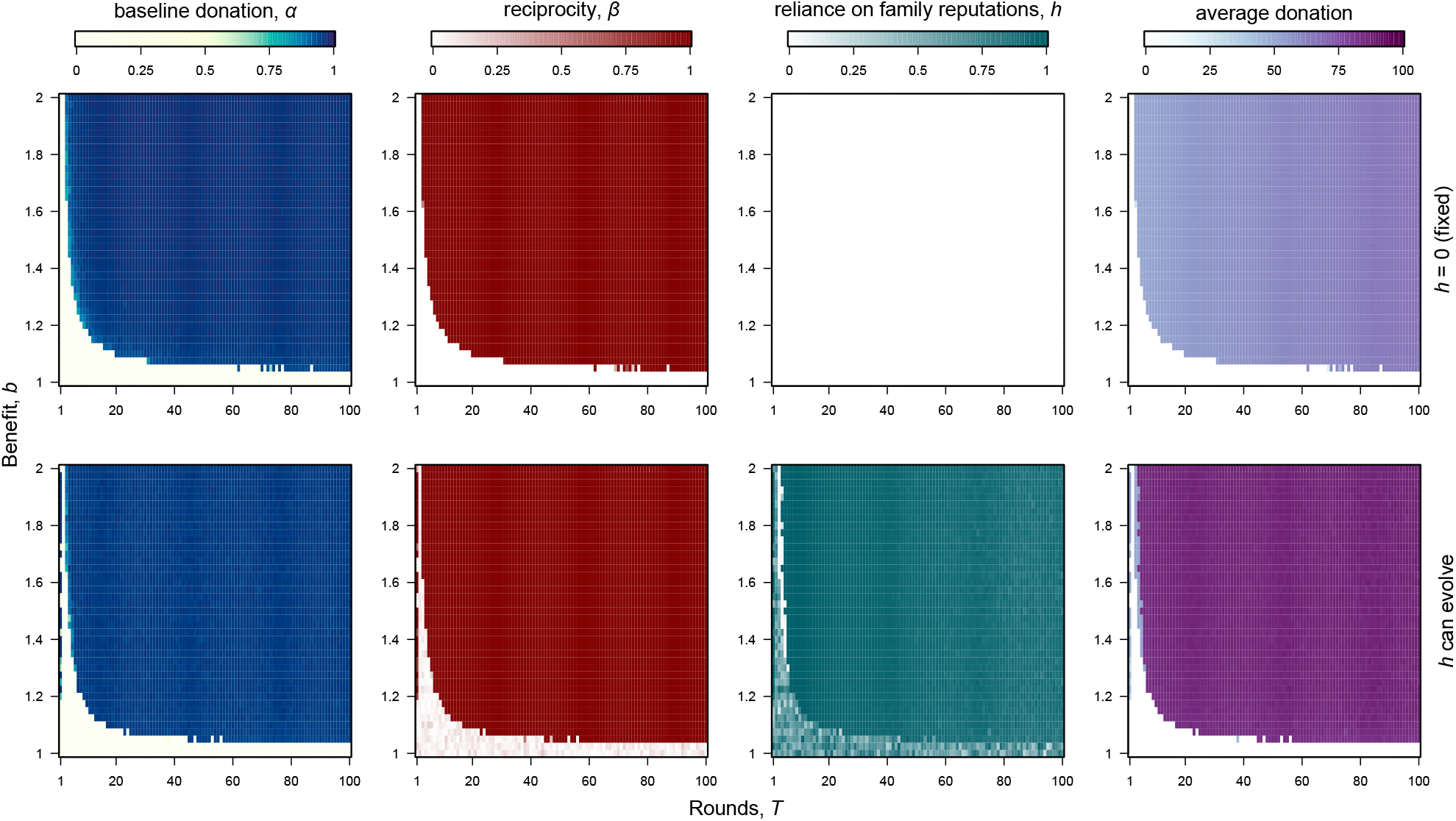
Evolution of reliance on family reputation with random initial reputations. Long-term trait averages across different numbers of rounds (*T*) and benefits (*b*), when individuals assign unknown partners a random initial reputation (*ρ*_ini_; see Section C.1 for simulation details). Reliance on family reputation (*h*) evolves in most of the parameters space, but does not expand the region where cooperation is favoured. Yet, when it evolves, *h* increases donations. Populations were initialised in the fully cooperative state without reputation inheritance (*α*= 1, *β*=0.99, *h* = 0). Reliance on family reputation *h* was either fixed at zero or allowed to evolve (top and bottom row, respectively). Parameters: *N* = 10^3^, *c* = 1, *q* = *s* = 1, ϵ _a_ =ϵ _e_ = 0, *ρ*_ini_ ∼ Unif[0, *d*_max_], *µ* = 10^−3^, *σ* = 0.1.

**Supplementary Figure 6.**
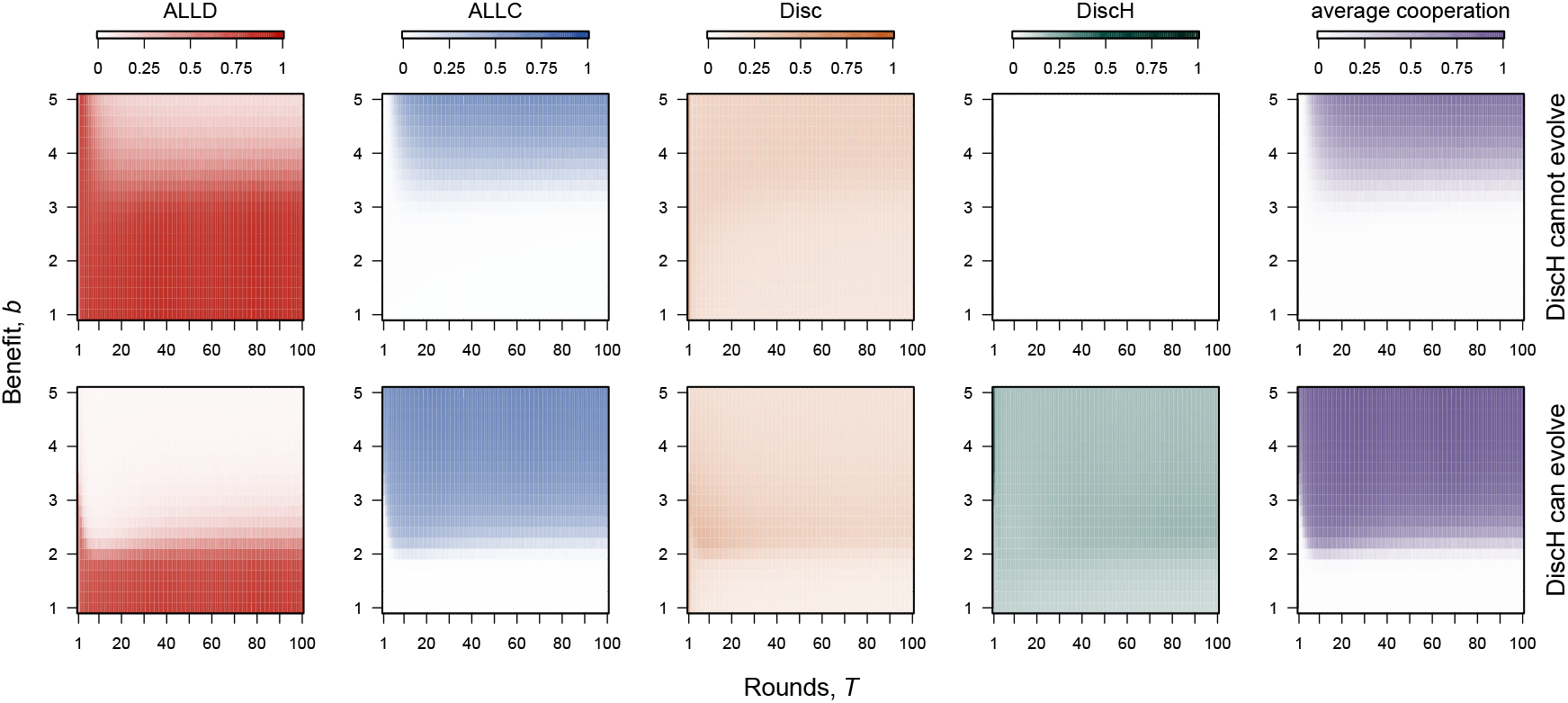
Classic pure-strategy indirect reciprocity game with image scoring. Long-term strategy and cooperation frequencies across different numbers of rounds (*T*) and benefits (*b*), when Discriminators who rely on family reputation (DiscH) are either not allowed or allowed to evolve (top and bottom row, respectively; see Section C.4 for simulation details). When allowed to evolve, DiscH increase the long-term average cooperation, especially under limited interactions and low benefits (low *T* and *b*, respectively). Unconditional defectors (ALLD) are mainly replaced by unconditional cooperators (ALLC), who are the most likely to inherit a GOOD reputation and benefit from it. Parameters: *N* = 10^3^, *c* = 1, *q* = *s* = 1, *ρ*_ini_ ∼ Unif(BAD, GOOD), ϵ _e_ =ϵ _a_ = 0.02, *µ* = 0.01.

**Supplementary Figure 7.**
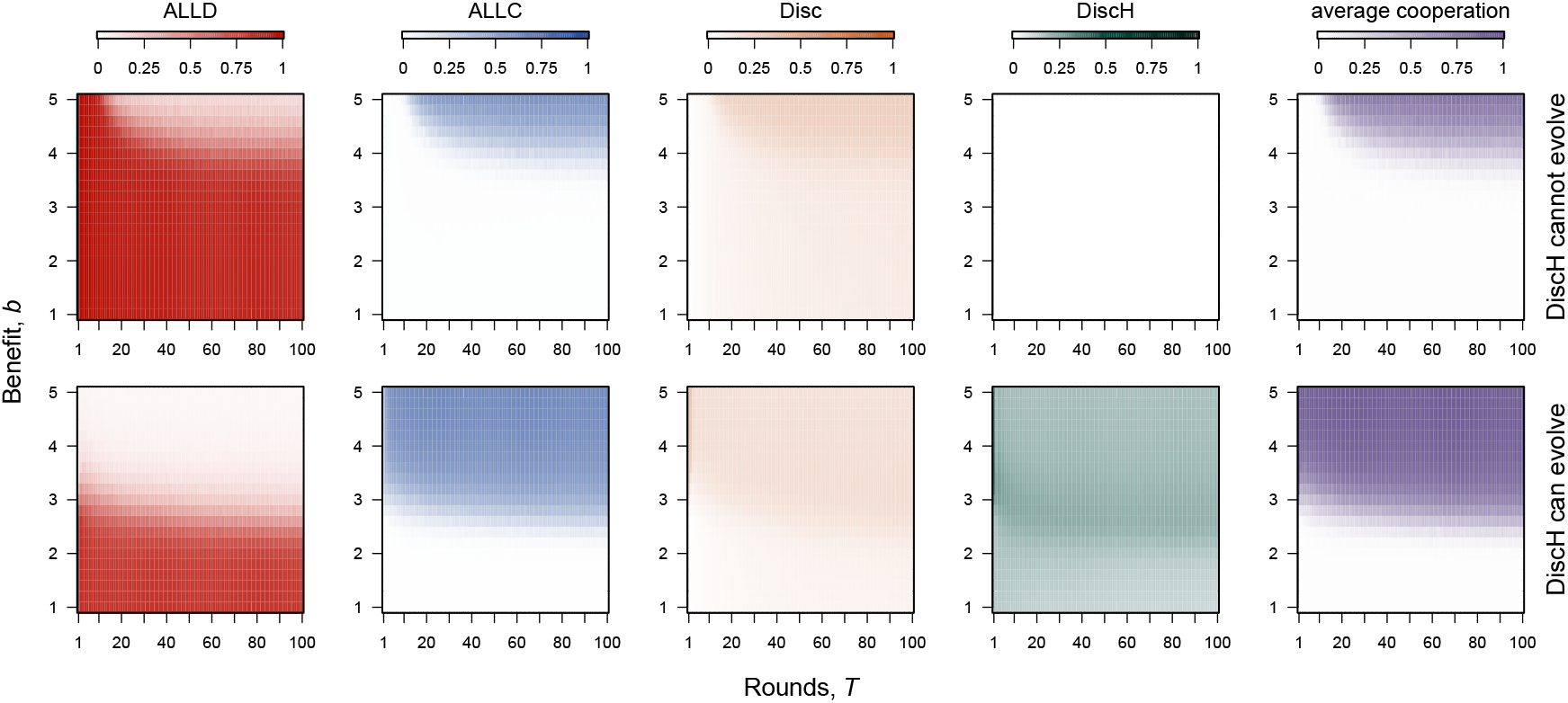
Classic pure-strategy indirect reciprocity game with trustful partners. Long-term strategy and cooperation frequencies across different numbers of rounds (*T*) and benefits (*b*), when Discriminators who rely on family reputation (DiscH) are either not allowed or allowed to evolve (top and bottom row, respectively; see Section C.4 for simulation details). Individuals are initially trustful, as they assign unknown partners a GOOD initial reputation. When allowed to evolve, DiscH again increase the long-term average cooperation, especially under limited interactions and low benefits (low *T* and *b*, respectively). Parameters: *N* = 10^3^, *c* = 1, *q* = *s* = 1, *ρ*_ini_ = GOOD, ϵ_e_ = ϵ_a_ = 0.02, *µ* = 0.01.

**Supplementary Figure 8.**
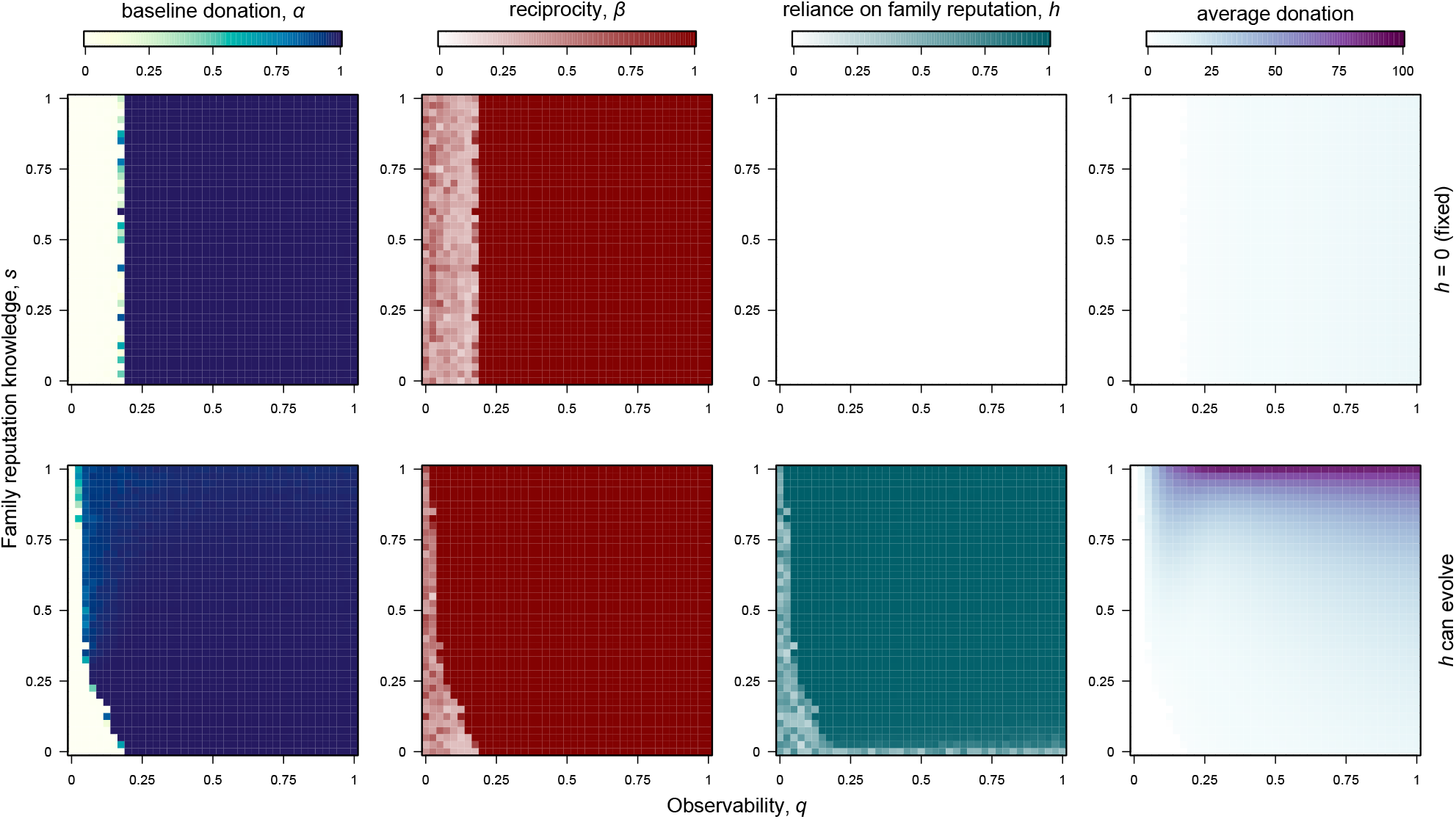
The evolution of reliance on family reputation compensates for low observability. Long-term trait averages for different levels of observability (*q*) and knowledge of family reputation (*s*). Reliance on family reputation stabilises co-operation, especially when observability is low (*q*<.2), and also improves donations. Lower knowledge of family reputation (*s*) disfavours reliance on family reputation. Populations were initialised in the fully cooperative state without reputation inheritance (*α*=1, *β*= 0.99, *h* =0). Reliance on family reputation *h* was either fixed at zero or allowed to evolve (top and bottom row, respectively; see Section C.2 for simulation details). Parameters: *N* = 10^3^, *c* = 1, *b* = 2, *T* = 20, ϵ_a_ =ϵ _e_ = 0, *ρ*_ini_ = 0, *µ* = 10^−3^, *σ* = 0.1.

**Supplementary Figure 9.**
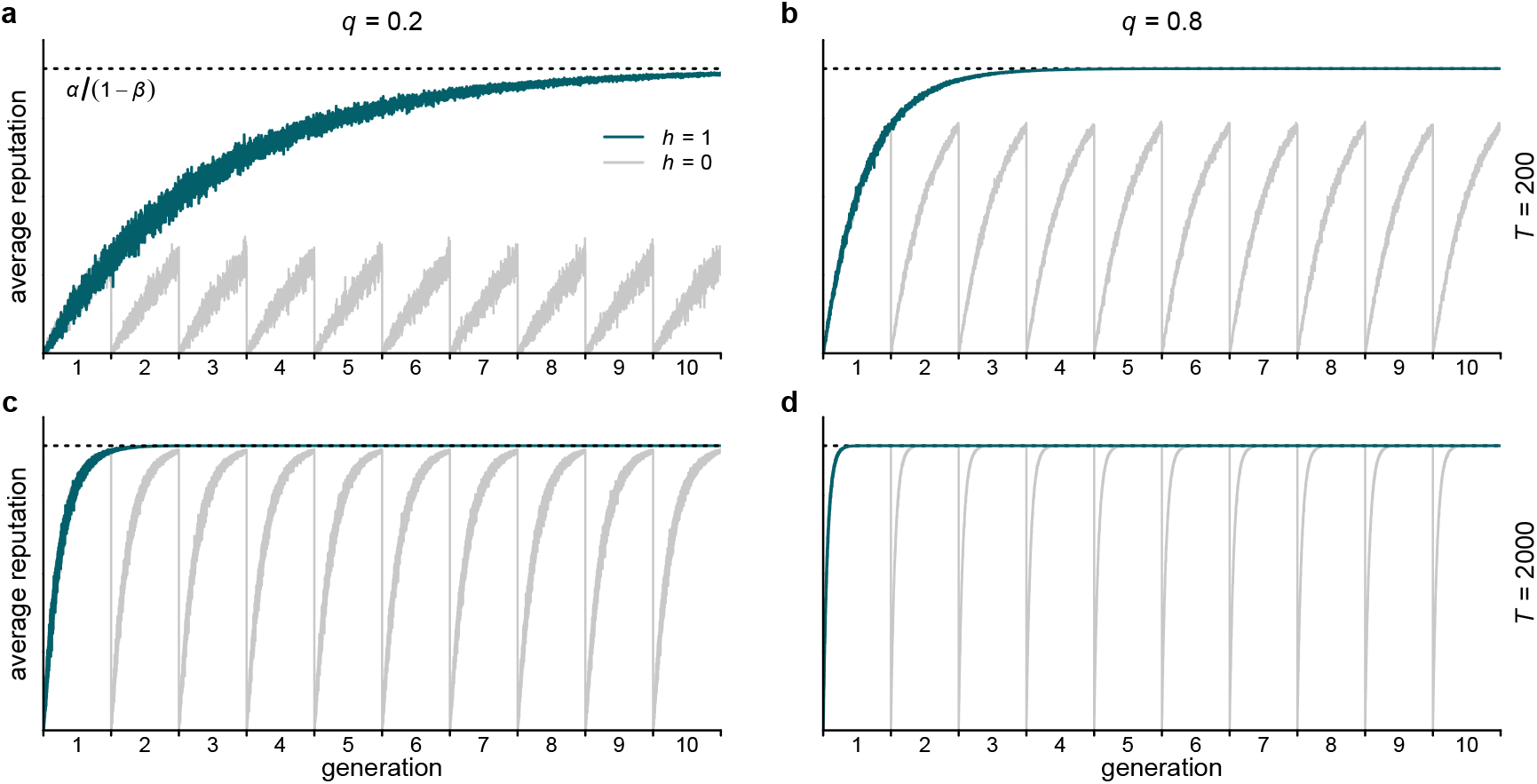
Reputation dynamics across generations. Average reputation in a fully cooperative monomorphic population (*α* = 1, *β*=0.99), under low and high observability (*q*=0.2, left panels, and *q*=0.8, right panels), and for different numbers of rounds *T* (top and bottom panels; see Section C.2 for simulation details). When family reputations are not inherited (*h*=0, light line), personal reputations reset every generation, and so, they can never reach equilibrium (dashed line) if observability (*q*) and the number rounds (*T*) are too low. When reputations are inherited (*h* =1, dark line), they build up cumulatively across generations, and rapidly reach equilibrium. Parameters: all panels, *N* = 10^3^, *c* = 1, *b* = 2, *s* = 1, ϵ_a_ = ϵ_e_ = 0, *ρ*_ini_ = 0, *µ* = *σ* = 0.

**Supplementary Figure 10.**
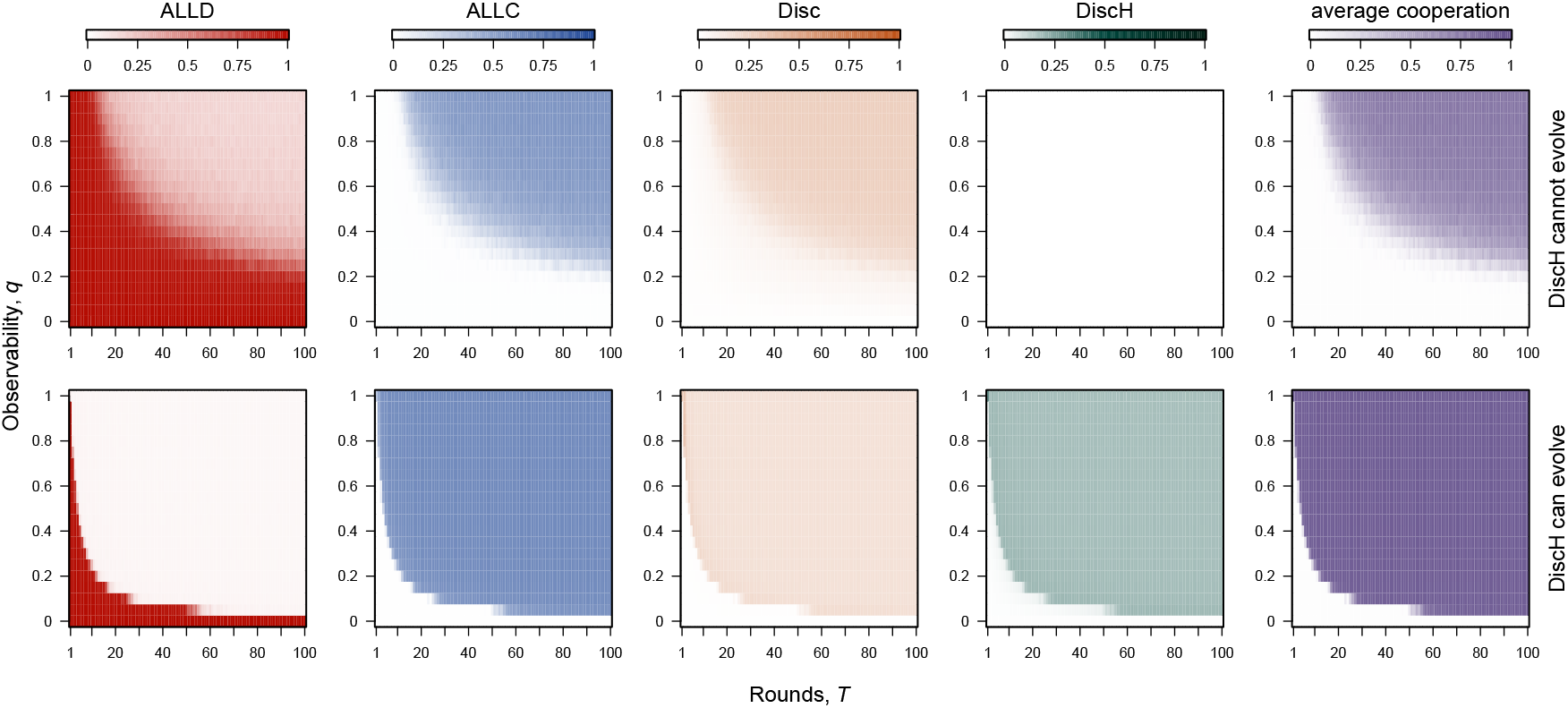
Imperfect observability in the classic pure-strategy indirect reciprocity game. Long-term strategy and cooperation frequencies across different numbers of rounds (*T*) and observability (*q*), when Discriminators who rely on family reputation (DiscH) are either not allowed or allowed to evolve (top and bottom row, respectively; see Section C.4 for simulation details). When allowed to evolve, DiscH increase the long-term average cooperation, even under low observability. Parameters: *N* = 10^3^, *c* = 1, *b* = 5, *s* = 1, *ρ*_ini_ = GOOD, ϵ_e_ = ϵ_a_ = 0.02, *µ* = 0.01.

**Table S1.**
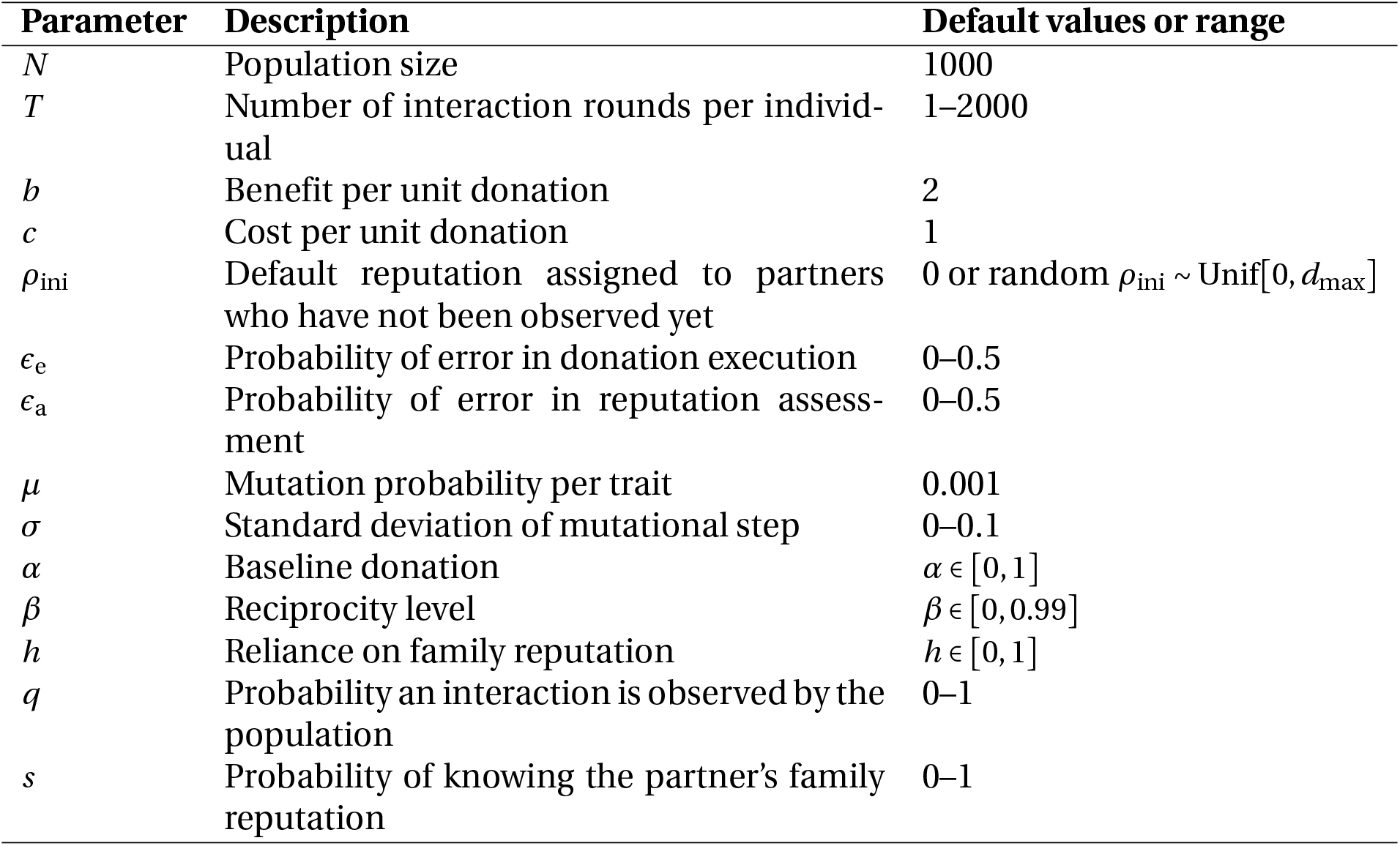
Simulation parameters.

| Parameter | Description | Default values or range |
| --- | --- | --- |
| $N$ | Population size | 1000 |
| $T$ | Number of interaction rounds per individual | 1–2000 |
| $b$ | Benefit per unit donation | 2 |
| $c$ | Cost per unit donation | 1 |
| $\rho_{\text{ini}}$ | Default reputation assigned to partners who have not been observed yet | 0 or random $\rho_{\text{ini}} \sim \text{Unif}[0, d_{\text{max}}]$ |
| $\epsilon_e$ | Probability of error in donation execution | 0–0.5 |
| $\epsilon_a$ | Probability of error in reputation assessment | 0–0.5 |
| $\mu$ | Mutation probability per trait | 0.001 |
| $\sigma$ | Standard deviation of mutational step | 0–0.1 |
| $\alpha$ | Baseline donation | $\alpha \in [0, 1]$ |
| $\beta$ | Reciprocity level | $\beta \in [0, 0.99]$ |
| $h$ | Reliance on family reputation | $h \in [0, 1]$ |
| $q$ | Probability an interaction is observed by the population | 0–1 |
| $s$ | Probability of knowing the partner's family reputation | 0–1 |

## Notes

### Competing Interest Statement

The authors have declared no competing interest.

